# Microsynteny Analysis Clarifies the Early Diversification of Cypriniform Fishes

**DOI:** 10.1101/2025.09.19.677351

**Authors:** Daniel J. MacGuigan, Hannah M. Waterman, Jessie A. Pelosi, Nathan J.C. Backenstose, Sarah L. Chang, Milton Tan, Fabrício Almeida-Silva, Trevor J. Krabbenhoft

## Abstract

While genomic data are powerfully informative for phylogenetic inference, traditional sequence-based analyses can be confounded by issues like homoplasy and incomplete lineage sorting, leaving deep nodes unresolved. Synteny, the conservation of gene order along a chromosome, presents a promising alternative source of phylogenetic information that is potentially less prone to these issues. Here we compare sequence and synteny phylogenetic phylogenomic approaches to unravel the evolutionary history of Cypriniformes, one of the most species-rich lineages of freshwater fishes. This clade has been plagued by uncertainty, particularly concerning the phylogenetic relationships between its four major subclades: Gyrinocheilidae (algae eaters), Catostomidae (suckers), Cobitoidei (loaches), and Cyprinoidei (minnows and carps). Phylogenomic approaches require high-quality genomic resources, which have been lacking for the family Gyrinocheilidae, a key lineage for understanding cypriniform evolution. To address this, we generated the first chromosome-level genome assembly and annotation for the Chinese Algae Eater (*Gyrinocheilus aymonieri*). We utilized this new genome along with 42 other fish genomes to conduct comprehensive phylogenomic analyses and compare results from sequence-based methods with a microsynteny-based approach. Although chromosome-level macrosynteny is broadly conserved among cypriniforms, we demonstrate that microsynteny has impressive power to resolve deep phylogenetic nodes. Both sequence-based and microsynteny-based analyses find that Gyrinocheilidae is the sister lineage to all other cypriniforms, resolving a long-standing phylogenetic point of contention. However, while sequence-based analyses provided weak support for the relationships among the remaining subclades, our microsynteny analysis revealed a novel and strongly supported sister relationship between Catostomidae and Cyprinoidei. This conclusion is supported by a 3- to 12-fold greater number of shared microsynteny clusters (synapomorphies) compared to the alternative topologies. We also demonstrate that microsynteny synapomorphies often arise from gene family expansions. Conservation of synteny concentrated in specific genomic regions may harbor the genomic underpinnings of the unique aspects of these clades and have played an important role in the evolutionary history of these taxa. Together, these results provide a robust phylogenomic backbone for Cypriniformes. More broadly, our findings highlight the power of microsynteny to resolve recalcitrant phylogenetic nodes where sequence data is uninformative, while also underscoring the critical importance of high-quality reference genomes, as our work also shows that including fragmented assemblies can lead to spurious results. Beyond phylogenomics, our results also provide a functional genomics glimpse into how conserved, clade-specific blocks of collinear genes (synteny synapomorphies) might contribute to what makes this hyperdiverse clade so unique and evolutionarily successful.

## INTRODUCTION

Genomic data provide a wealth of information for phylogenetic inference, but most genome-scale analyses focus on nucleotide characters from sequence alignments. For example, many studies have employed genomic reduced representation or marker panel approaches (RAD-seq, transcriptome sequencing, ultra-conserved elements, exon capture, etc.) and, increasingly, whole genome alignments (Jarvis et al. 2014; Armstrong et al. 2020) or gene-trees derived from orthology analysis (Waterhouse et al. 2018; Manni et al. 2021; Tan et al. 2021). While nucleotide and protein sequences are powerfully informative for phylogenetic analyses, their utility in unraveling complex phylogenetic patterns and more recalcitrant regions of evolutionary history can be limited by homoplasy and phylogenetic signal saturation. The small numbers of potential character states coupled with high substitution rates can cause issues for phylogenetic inference, especially in ancient rapid radiations (Whitfield and Lockhart 2007).

Crucial information about evolutionary relationships can also be embedded within genome architecture. While using gene content and order (i.e., synteny) to understand phylogenetic relationships is not new (Sturtevant and Dobzhansky 1936; Dobzhansky and Sturtevant 1938), advances in genome sequencing, assembly, and annotation have made this approach tractable for a broad diversity of species (Steenwyk and King 2024a). In this context researchers often differentiate between macrosynteny (large-scale chromosomal rearrangements such as fusions or fissions) and microsynteny (local gene collinearity within chromosomes) (Yu et al. 2024). The utility of both macrosynteny (Parey et al. 2023) and microsynteny patterns, especially gene collinearity (Zhao et al. 2021), as phylogenetically informative data sources has recently been demonstrated for resolving longstanding phylogenetic challenges (Schultz et al. 2023). In some cases, phylogenetic signal is congruent between sequence-based and synteny-based approaches (Parey et al. 2023). However, the potential for homoplasy may be lower for synteny-based phylogenetic methods compared to nucleotides; the large numbers of possible character states means that convergent rearrangements of gene collinearity are highly unlikely to occur by chance (Rokas and Holland 2000; but see MacGuigan et al. 2023). Likewise, synteny methods can be more robust to the confounding effects of gene flow (Ding et al. 2023). Thus, in some situations, synteny may be more informative about evolutionary history than sequence data, such as in cases involving long branch attraction or rapid bursts of speciation, where incomplete lineage sorting is likely to occur. However, caution is still warranted in the interpretation of apparent phylogenetic signal in synteny data; hybridization, incomplete lineage sorting, and selection favoring parallelism can contribute to misleading results (Steenwyk and King 2024a). Few studies have explicitly compared the phylogenetic signal in sequences versus synteny (but see Zhao et al. 2021), and to our knowledge there have been no explicit topology tests using microsynteny data.

To compare the phylogenetic power of synteny-based and sequence-based approaches, we examined the evolutionary history of Cypriniformes. Cypriniformes is a clade that encompasses 4,908 species of fish (Fricke et al. 2025), including some of the most economically important freshwater fisheries on the planet. For example, the cypriniform species Common Carp (*Cyprinus carpio*), Silver Carp (*Hypothalmichthys molitrix*), and Grass Carp (*Ctenopharyngodon idella*) had a combined 15 million tons of fish harvested in 2022 (FAO 2024). Cypriniform fishes made up 18% of global aquatic animal production for fisheries and aquaculture in 2022 (FAO 2024). In addition to their role in global food production, Cypriniformes also includes the biomedically-important zebrafish (*Danio rerio*), a vertebrate model for developmental and neurobiology and one of the first vertebrate genomes to be sequenced (Mayden et al. 2007; Howe et al. 2013).

Cypriniformes is comprised of four major subclades: Gyrinocheilidae (algae eaters; 3 species), Catostomidae (suckers; 86 species), Cobitoidei (loaches; 1,429 species), and Cyprinoidei (minnows and carps; 3,390 species) (Tan and Armbruster 2018; Fricke et al. 2025 http://researcharchive.calacademy.org/research/ichthyology/catalog/SpeciesByFamily.asp). Numerous studies have presented phylogenetic hypotheses for the evolutionary relationships within Cypriniformes based on both morphological (e.g. Conway et al. 2010; Conway 2011) and molecular datasets (Saitoh et al. 2006; Mayden et al. 2008; Stout et al. 2016; Hirt et al. 2017; Tao et al. 2019). However, there remains little consensus about the deepest relationships among the four major subclades of Cypriniformes (Fig. 1A,B), with some authors altogether questioning the validity of molecular-based phylogenetic hypotheses (Britz and Conway 2011). The largest molecular phylogenetic dataset for cypriniforms to date contained 219 anchored hybrid enrichment loci sampled for 172 taxa. Phylogenetic analyses of these data produced three different topologies (all strongly supported) for relationships among the four main cypriniform subclades, depending on inference method employed (Fig. 1B, Stout et al. 2016). This uncertainty likely stems from few informative characters due to rapid early diversification leading to the major extant cypriniform clades.

**Figure 1.**
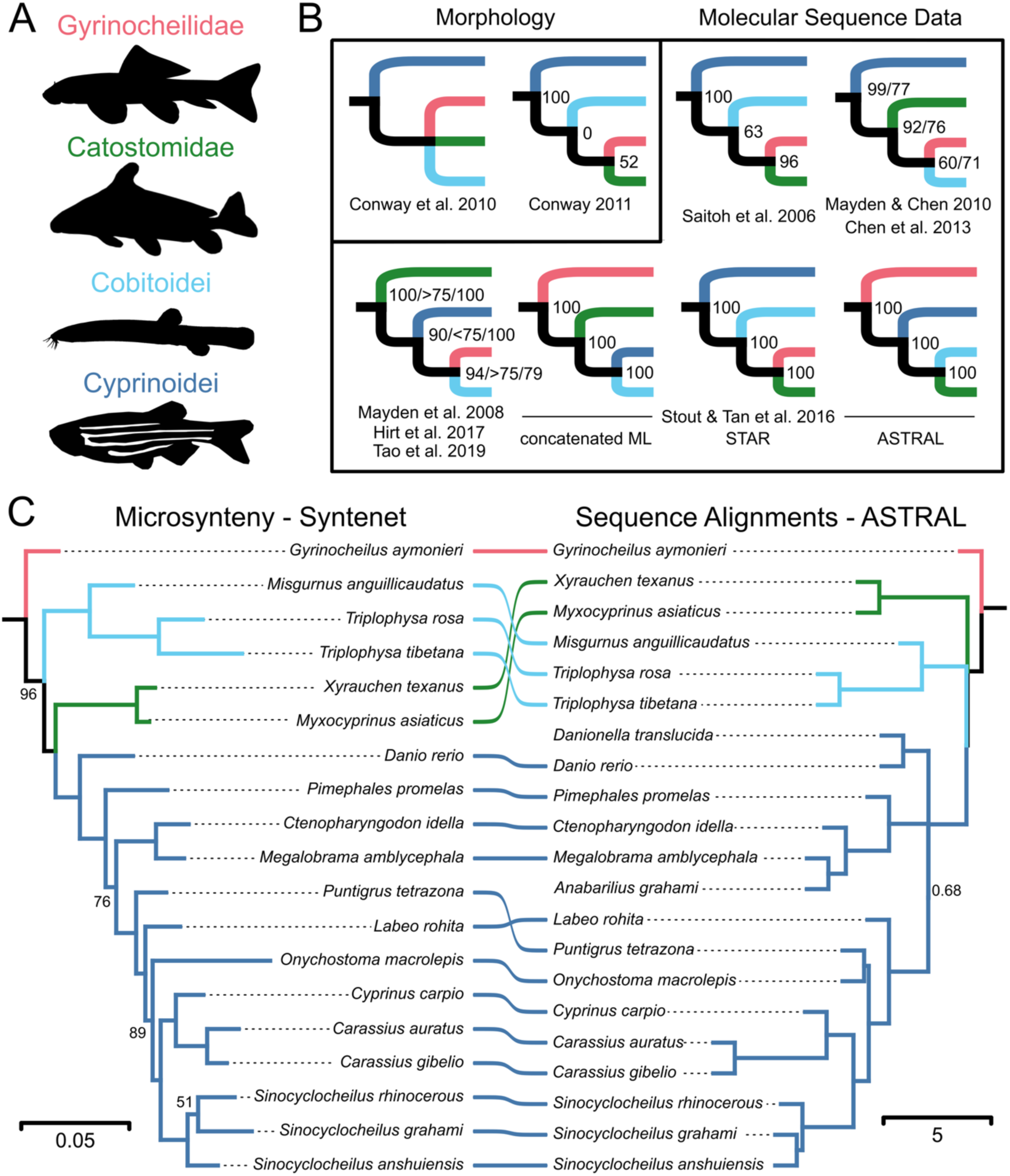
Sequence and synteny phylogenies for Cypriniformes. A) The four main lineages of Cypriniformes. B) Summary of prior morphological and molecular phylogenetic hypotheses for Cypriniformes. C) Microsynteny (51,040 clusters) and sequence alignment (8,731 orthogroups) phylogenies. Outgroup taxa have been pruned. Support values less than 100% bootstrap support (left) or less than 1.0 local posterior probability (right) are hidden. Branch lengths are in units of expected number of changes per synteny cluster (left) and coalescence units (right). Note that ASTRAL does not estimate terminal branch lengths. Fish silhouettes were obtained from PhyloPic.org.

**Table 1.** *Gyrinocheilus aymonieri* genome assembly statistics.

|  |  |  |
| --- | --- | --- |
| Assembly | Number of contigs | 387 |
|  | Longest contig (Mb) | 10.18 |
|  | Contig N50 (Mb) | 2.92 |
|  | Scaffolded assembly length (Gb) | 0.458 |
|  | Number of scaffolds | 288 |
|  | Scaffold N50 (Mb) | 18.57 |
|  | N (%) | 0.01 |
|  | GC (%) | 39.82 |
| BUSCO v5<br>(actinopterygii_odb10) | Complete (%) | 97.5 |
|  | Complete single copy (%) | 96.3 |
|  | Complete duplicated (%) | 1.2 |
|  | Fragmented (%) | 1.0 |
|  | Missing (%) | 1.5 |
| Repetitive elements | Total (%) | 23.06 |
|  | SINEs (%) | 0.08 |
|  | LINEs (%) | 1.42 |
|  | LTR elements (%) | 2.33 |
|  | DNA transposons (%) | 2.28 |
|  | Unclassified (%) | 14.03 |
| Annotation | Predicted genes | 27,956 |
|  | Predicted transcripts | 27,956 |
|  | Mean [median] gene length (bp) | 7,608.0 [3,512.5] |
|  | Mean [median] exon length (bp) | 177.9 [124] |
|  | Mean intron length (bp) | 710.2 [213] |
|  | Mean [median] exons per transcript | 9.4 [6] |
|  | Mean [median] introns per transcript | 8.4 [6] |

Phylogenetic analyses of DNA sequence data alone may be insufficient to resolve the deepest nodes within Cypriniformes. Fortunately, this clade is ideally suited for alternative phylogenomic approaches, such as analyses of microsynteny. Recent sequencing projects have resulted in chromosome-level annotated reference genomes for at least two representatives of Catostomidae, Cobitoidei, and Cyprinoidei. However, there are currently no genomes available for the fourth major cypriniform clade, Gyrinocheilidae. This important lineage is currently represented by one genus, *Gyrinocheilus*, with only three recognized species (Tan and Armbruster 2018).

Here, we present a high quality, chromosome-level genome assembly of the Chinese Algae Eater (*Gyrinocheilus aymonieri*). We reconstructed sequence-based phylogenetic trees, performed synteny-based phylogenetic analyses, investigated clade-specific microsyntenic patterns, and performed macrosynteny analysis to evaluate alternative topologies within the Cypriniformes clade. We explored the application of micro- and macrosynteny information for evaluating alternative topologies in cases where there is insufficient phylogenetic signal in sequence data, or when the signal is ambiguous and potentially misleading. Using this combined approach, we find strong support for a novel phylogenetic hypothesis of cypriniform relationships and provide an overview of the application and caveats associated with microsynteny-based phylogenies toward complex phylogenetic patterns.

## MATERIALS AND METHODS

### Genome sequencing, assembly, and scaffolding

A single adult *Gyrinocheilus aymonieri* individual was purchased from an aquarium fish store (Pets Plus, Lockport, New York, USA), sacrificed with a lethal dose of MS-222 (>400 mg/L) (IACUC protocol #201900161), and dissected immediately. Muscle and gill tissues were removed and placed on dry ice and stored frozen at -80°C. High molecular weight DNA was extracted from muscle and gill tissues using a 500G Genomic Tip (Qiagen, Inc., Germantown, Maryland, USA), following the manufacturer’s protocol. Additional DNA was extracted from muscle tissue with an Invitrogen Life Technologies (Carlsbad, California, USA) PureLink Genomic DNA Mini Kit for short read sequencing. DNA was quantified with Qubit v4 dsDNA Broad Range kit (Thermo Fisher Scientific, Waltham, MA, USA).

Two high molecular weight DNA libraries were prepared (one from muscle and one from gill tissue) with ligation kits (LSK-109) from Oxford Nanopore Technologies (ONT; Oxford, UK). Each library was run on an ONT MinION flow cell (R9.4.1) on the GridION platform. Nanopore reads were basecalled in high-accuracy mode with Guppy v3.0.6. To generate additional high-accuracy short reads, DNA extracted with the PureLink DNA Mini Kit was sent to Novogene, Inc (Davis, California, USA) for library prep and sequencing on an Illumina Novaseq6000 with a PE150 layout.

For genome assembly, Minimap2 v2.17 (Li 2018) was used to perform all-by-all mapping of the long reads with the *–x ava-ont* flag. The resulting pairwise alignment file and raw reads were assembled using Miniasm v0.3 (Li 2016). Two iterations of long read polishing were completed by mapping ONT reads to the assembly with Minimap2, and a consensus sequence was generated with racon v1.4.11 (Vaser et al. 2017). Illumina reads were processed by removing adapter sequence, matching proper pairs, and filtering for quality with trim_galore v0.6.10 *(--paired, --quality 20*; Martin 2011) and used in two rounds of polishing with Pilon v1.23 (Walker et al. 2014). For polishing, the trimmed Illumina reads were mapped to the racon-polished assembly using BWA-MEM v0.7.17 (Li and Durbin 2009; Li 2013). The mapped reads were filtered for minimum mapping quality score (-*q 15*) using samtools v1.15 (Li et al. 2009) and used in Pilon to correct for variation within the assembly (*--frags, --fix-all*).

Purge_haplotigs v1.0.4 (Roach et al. 2018) was used to identify and remove possible haplotigs and collapsed repeats, using *-l 5*, *-m 20*, and *-h 65*, for low, middle, and high coverage, thresholds, respectively. Microbial and other contamination was identified in the final assembly using Kraken2 v2.1.0 (Wood et al. 2019) with the preassembled databases archaea, bacteria, plasmid, viral, human, fungi, plant, protozoa, as well as NCBI’s nr/nt databases, UniVec, and UniVec_Core. A Hi-C library from skeletal muscle tissue was generated and sequenced on an Illumina NovaSeq 6000 by Phase Genomics Inc. (Seattle, WA, USA) for genome scaffolding. The contig-level assembly was scaffolded using Juicer v1.6 (Durand et al. 2016) with default parameters with the enzyme *DpnII* (GATCGATC) and then piped into 3D-DNA v180922 (Dudchenko et al. 2017) using the following options: editor-coarse-stringency, polisher-coarse-stringency, and *splitter-coarse-stringency = 10*, *editor-fine-resolution*, *polisher-fine-resolution*, and *splitter-fine-resolution = 25,000*, and editor-coarse-resolution, polisher-coarse-resolution, and *splitter-coarse-resolution = 100,000*. BUSCO v5.1.2 (Seppey et al. 2019) was used to assess the completeness of the final assembly (without gene models) using the actinopterygii_odb10 core gene set.

### Genome annotation

We used a combination of the BRAKER v3.0.2 pipeline (Gabriel et al. 2024) and GeMoMa v1.9 (Keilwagen et al. 2019) to annotate the scaffolded *Gyrinocheilus aymonieri* genome. First, we generated a custom repeat library for *G. aymonieri* using six rounds of RepeatModeler v.2.0.1, including the optional LTR structural discovery pipeline (Ellinghaus et al. 2008; Ou and Jiang 2018; Flynn et al. 2020). We filtered the custom repeat library to remove protein-coding coding genes retrieved from the UniProt Swiss-Prot database (http://www.uniprot.org/; last accessed March 1, 2023). We then generated a repeat GFF file using RepeatMasker v.4.1.1 (Smit AFA, Hubley R, Green P. 2015. RepeatMasker open-4.0. Available from: http://www.repeatmasker.org) with the filtered *G. aymonieri* repeat library and the Dfam v.3.2 repeat database (Storer et al. 2021). Bedtools v.2.30.0 was used to generate as soft-masked genome assembly file (Quinlan and Hall 2010).

Both RNA-seq data and protein sequences were used for the BRAKER3 annotation pipeline. We retrieved 8.7 Gbp of paired-end Illumina RNA-seq data for *G. aymonieri* from the NCBI SRA (SRX2479405 and SRX3153282). Raw RNA-seq data were aligned to the masked genome using HISAT2 v.2.2.1 with default parameters (Kim et al. 2019). We used two sets of protein evidence for BRAKER3. First, we downloaded predicted proteins from the NCBI RefSeq database for six cypriniform species: *Cyprinus carpio* (GCF_000951615.1), *Myxocyprinus asiaticus* (GCF_019703515.2), *Ctenopharyngodon idella* (GCF_019924925.1), *Labeo rohita* (GCF_022985175.1), *Carassius gibelio* (GCF_023724105.1), and *Xyrauchen texanus* (GCF_025860055.1). Additionally, we downloaded the metazoan OrthoDB v.11 protein database (Kuznetsov et al. 2023). The RNA-seq alignments and protein data were used to run BRAKER3 with default parameters. AUGUSTUS (Stanke et al. 2008) and GeneMark-ETP (Brůna et al. 2024) hints were then combined using TSEBRA v.1.1.0 with the default parameters for BRAKER3 (Gabriel et al. 2021).

In addition to BRAKER3, we used the homology-based pipeline GeMoMa to generate a set of gene predictions for *G. aymonieri*. Using the masked genome and default GeMoMa parameter settings, we combined separate sets of prediction using NCBI RefSeq protein evidence from 15 cypriniform genome assemblies: *Danio rerio* (GCF_000002035.6), *Cyprinus carpio* (GCF_018340385.1), *Sinocyclocheilus anshuiensis* (GCF_001515605.1), *S. rhinocerous* (GCF_001515625.1), *S. grahami* (GCF_001515645.1), *Carassius auratus* (GCF_003368295.1), *Pimephales promelas* (GCF_016745375.1), *Megalobrama amblycephala* (GCF_018812025.1), *Puntigrus tetrazona* (GCF_018831695.1), and the six species used for BRAKER3.

We combined BRAKER3 and GeMoMa predictions using EVidenceModeler v2.0.0 (Haas et al. 2008). We converted the BRAKER3 and GeMoMa predictions using the appropriate EvmUtils scripts and merged annotations with the following parameters: weight for BRAKER3 *predictions = 1*, weight for GeMoMa *predictions = 3*, *--segmentSize 1000000*, *--overlapSize 100000*. Annotation statistics were calculated using a custom R script and the web tool gVolante v.2.0.0 was used to assess annotation quality and completeness by querying the BUSCO v5 actinopterygii_odb10 ortholog dataset (Nishimura et al. 2017). Additionally, we compared the completeness and phylogenetic consistency of our *G. aymonieri* predicted proteins to other taxa using OMArk (Nevers et al. 2025).

### Sequence-based Phylogenetic Analyses

To examine the phylogenetic placement of Gyrinocheilidae, we downloaded 42 additional fish genome assemblies and their annotations from NCBI (Table S1). These included five species of Clupeiformes as outgroups and representatives from all major clades of the Ostariophysi: Gonorynchiformes (milkfish and allies), Gymnotiformes (knifefishes), Siluriformes (catfishes), and Cypriniformes (minnows, suckers and loaches). Thirty three of these genomes were chromosome-level assemblies, nine were scaffold-level, and one was contig-level. All genome annotations were filtered to retain only the longest isoforms using AGAT v1.1.0 (Dainat et al. 2025).

For our sequence-based phylogenetic inference, we used OrthoFinder v.2.5.4 to identify sets of single-copy orthologous genes (Emms and Kelly 2019). We then performed three phylogenetic analyses. All analyses utilized MAFFT v7.490 (Katoh and Standley 2013) to generate alignments of amino acid sequences. First, we used 423 single-copy orthogroups (with a minimum of 87.2% of species having single-copy genes in any orthogroup) to build a species tree with OrthoFinder’s STAG consensus species tree approach (Emms and Kelly 2018). Next, we concatenated the 423 single-copy orthogroup alignments and performed maximum likelihood phylogenetic inference with IQTree v.2.2.0 (Minh et al. 2020b), selecting the best-fit substitution model with ModelFindera (*-m MFP*) (Kalyaanamoorthy et al. 2017) and performing 1,000 ultrafast bootstrap replicates (*-B 1000*) (Hoang et al. 2018).

Given that out dataset contains several known (paleo)tetraploid taxa (Table S1), we used custom bash scripts to identify orthogroups that were single-copy for diploid taxa but contained up to two gene copies for “polyploid” (but functionally diploid) taxa. This custom filtering allowed us to recover several thousand orthogroups, far more than were recovered with strict single-copy filtering. We applied an additional set of filtering across a range of missing taxa values, from 0% missing taxa to 20% missing taxa (Table S2). Since we observed a drop-off in the number of loci retained above the 9% (39/43 taxa) cutoff, we selected this as our threshold for retaining alignments. The resulting 8,731 multi-copy orthogroups were used to perform a coalescent-based species tree analysis with ASTRAL-Pro v.1.15.1.3 (Zhang et al. 2020), a modification of the original ASTRAL algorithm (Mirarab et al. 2014) that accommodates paralogy. Input gene trees for ASTRAL-Pro were inferred using IQTree, with ModelFinder deployed to identify the best-fit model (*-m MFP*). Default settings were used for ASTRAL-Pro.

In addition to these phylogenetic inference approaches, we performed gene and site concordance factor analyses using IQTree. Gene trees for the 423 single-copy orthogroups were inferred using IQTree. The concatenated IQTree topology was used as the guide tree when calculating concordance factors (Minh et al. 2020a). For calculating site concordance factors, we randomly sampled 100 quartets around each internal branch (*--scf 100*).

### Synteny-based Phylogenetic Analyses

After preliminary synteny analyses, we excluded four taxa due to issues with annotation or assembly quality (Fig. S3), resulting in a dataset of 39 fish genomes, including our chromosome-level *Gyrinocheilus* assembly. To explore broad patterns of macrosynteny among otocephalans, we used the GENESPACE (v.1.2.3) pipeline (Lovell et al. 2022), which incorporates MCScanX (Wang et al. 2012), to infer syntenic orthologues and paralogues among the 33 species with chromosome-level assemblies. Since our dataset contained several polyploid taxa, we used a custom bash script to filter the DIAMOND results for paralogs by retaining only the top hits based on percent sequence identity (Buchfink et al. 2021) prior to runningGENESPACE. We used a custom bash script to order and reorient chromosomes for the GENESPACE synteny riparian plot.

We used the Syntenet v1.3.0 R package (Almeida-Silva et al. 2023) to infer phylogenetic relationships from patterns of microsynteny. Prior to running the Syntenet analyses, we use a custom bash script to filter the pairwise DIAMOND results, retaining only hits with sequence length >30 amino acids and >70% sequence identity. Briefly, Syntenet identifies syntenic gene pairs between species using the MCScanX algorithm, constructs a network of syntenic links across all species, and performs clustering analyses to identify synteny clusters in the network. These synteny clusters were then converted to binary presence-absence characters for phylogenetic analysis. The Syntenet *infer_microsynteny_phylogeny* function was used to reconstruct the phylogeny from the binary synteny matrix; this function follows (Zhao et al. 2021) in using IQTree with a Jukes-Cantor type model using free and equal rates and state frequencies by maximum-likelihood (*-m Mk+R+FO*) and 1000 ultrafast bootstrap replicates. To visualize the synteny clusters as a heatmap, we reduced noise by excluding clusters present in only one taxon or shared by more than 90% of taxa.

### Genomic Distribution of Microsynteny Phylogenetic Signal

To compare the strength of support for the phylogenetic resolution of Catostomidae, Cobitoidei, and Cyprinoidei, we used the Syntenet “find_GS_clusters” function, which calculates the number of clade-specific synteny clusters (hereafter referred to as synteny synapomorphies). We calculated synteny synapomorphies for the three possible topologies using a range of “min_percentage” values (the minimum percentage of species required to consider a cluster “clade-specific”). We visualized the genomic distribution of phylogenetically informative synteny clusters using the R package RIdeogram v.0.2.2 (Hao et al. 2020).

We examined the genomic distribution of microsynteny phylogenetic signal for the clade of Catostomidae plus Cyprinoidei. Syntenic gene density was calculated as the number of genes which are synteny synapomorphies, normalized by the total number of genes in each window. A syntenic gene density of 1 indicates that all genes in the window are synteny synapomorphies. We calculated this metric in sliding windows across the *Danio rerio* genome using a window size of 3 Mbp and a slide size of 0.5 Mbp. *Danio rerio* was selected as the representative taxon due to its importance as a model organism and long history of genomic study. We used a custom R script to identify and visualize synteny clusters in the three regions of the *Danio rerio* genomic regions with the highest syntenic gene density. Gene products and functional information for the genes comprising each cluster were extracted from the NCBI annotations.

To broadly examine patterns of synteny phylogenetic information content, we also calculated the genomic distribution of clade-specific synteny cluster for five clades of varying evolutionary ages: Ostariophysi (∼154-170 Ma, Hughes et al. 2018), Cypriniformes (∼85-115 Ma, Hughes et al. 2018), Cyprinoidei (∼67-98 Ma, Hirt et al. 2017), Catostomidae (72-105 Ma, Hirt et al. 2017), and Cobitoidea (∼50-78 Ma, Hughes et al. 2018). For each clade, we used the Syntenet “find_GS_clusters” function, specifying “min_percentage = 100” to identify synteny clusters that were shared by all members of the clade. Synteny synapomorphies for each clade were visualized for a representative taxon using the sliding window approach described above. Additionally, we visualized the distribution of normalized syntenic gene density using heatmaps of 1 Mbp genomic windows.

## RESULTS

### Gyrinocheilus aymonieri Genome Assembly and Annotation

We generated 2,949,777 Oxford Nanopore Technologies reads for *Gyrinocheilus aymonieri*, comprising 17.4 Gbp for a total average coverage of 38x coverage (read N50 = 11.8 Kbp [Supplemental Table 1]). Reads derived from gill and muscle tissue DNA represented 17.5x and 20.5x coverage, respectively. An additional 30.25 Gbp of Illumina sequence data (66x coverage) were used to polish the initial long-read assembly (Supplementary Table 1). The Hi-C dataset comprised another 80.2 Gbp (174x coverage) of Illumina sequence data for genome scaffolding.

The initial *G. aymonieri* assembly was highly contiguous, containing 387 contigs with an N50 of 2.92 Mbp. Scaffolding with Hi-C data produced 288 total scaffolds with an N50 of 18.6 Mb (Fig. S1). The 24 largest scaffolds (hereafter referred to as chromosomes) comprised 98% of the total assembly length. The genome of *G. aymonieri* was notable for having a smaller genome size (0.46 Gbp) than any of the cypriniforms included in this study; the cypriniform species with the next smallest genome assembly is the loach *Triplophysa tibetana* (0.65 Gbp) and the average genome size of the diploid taxa was 0.95 Gbp.

Functional annotation of the scaffolded *G. aymonieri* assembly identified 23.1% of the genome as repetitive elements and produced a total of 27,956 predicted gene models. The predicted genes contained 98.4% complete (96.3% single-copy) and 0.4% fragmented BUSCO genes (Fig. S1). OMArk analyses of the 16,357 hierarchical orthologous groups (HOGs) for Otophysi found 97.6% were complete (93.4% single-copy) and 2.4% were missing in the *G. aymonieri* predicted proteins (Fig. S1). OMArk further showed that 90.0% of the *G. aymonieri* protein sequences had phylogenetically consistent lineage placement.

### Macrosynteny patterns

We observed strongly conserved macrosynteny among cypriniforms, with a clear 1:1 chromosome relationship (or 1:2 relationship for polyploid taxa) between *G. aymonieri* and other cypriniform species (Fig. 2B). There are no major chromosomal rearrangements among the cypriniform species examined, a lineage-specific fusion in *Gyrinocheilus* and an independent chromosomal fusion in Cyprinoidei shared by *Megalobrama amblycephala* (Wuchang Bream) and *Ctenopharyngodon idella* (Grass Carp) (Fig. 2B-V); these latter two species are the only xenocyprinids included in the present analysis, so this shared macrosynteny rearrangement is likely phylogenetically informative.

**Figure 2.**
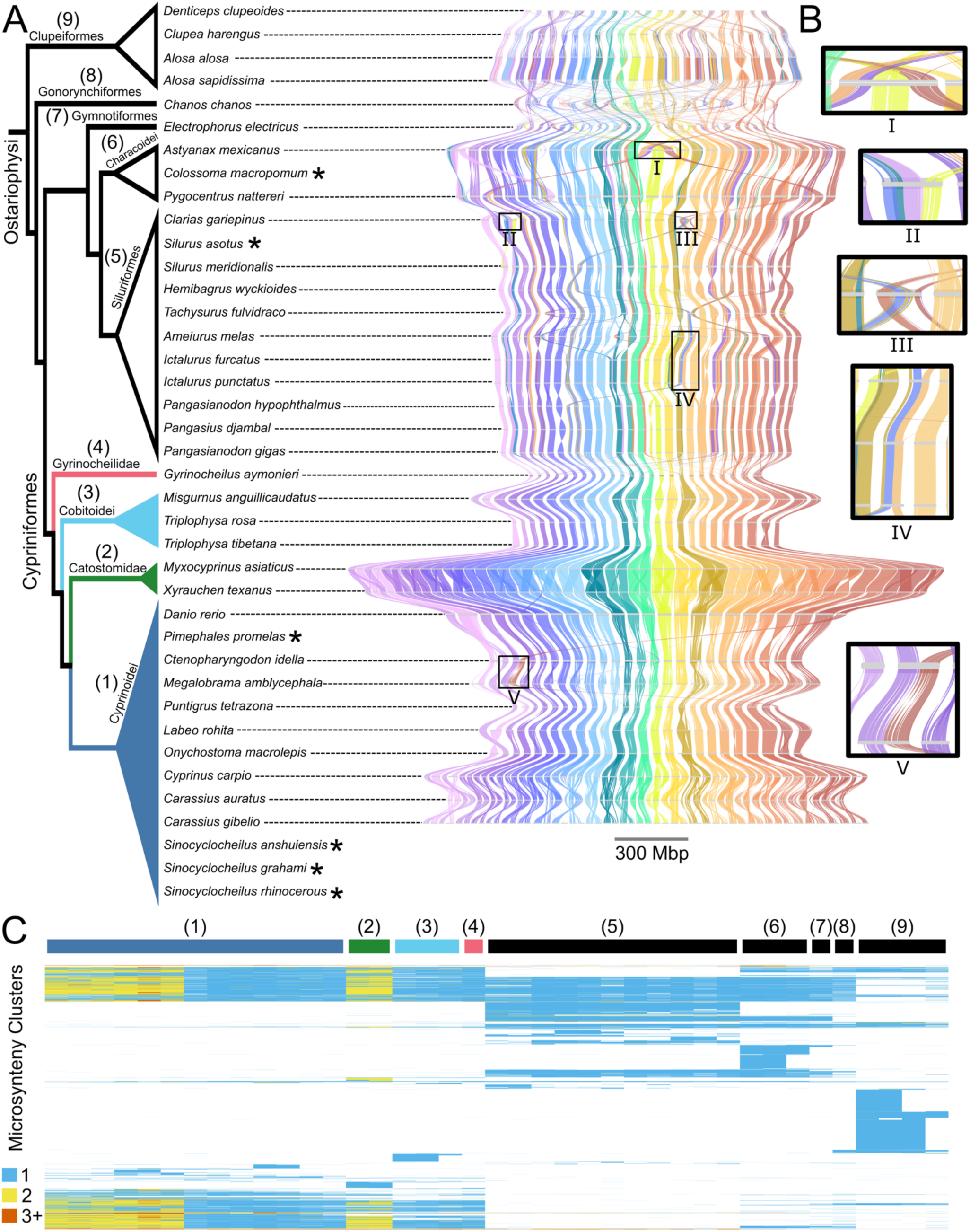
Phylogenetic informativeness of macrosynteny versus microsynteny. A) Microsynteny-based cladogram of Ostariophysi. Asterisks indicate taxa omitted from the GENESPACE riparian plot of macrosynteny in B) Colors represent blocks of conserved synteny between species. Chromosomes are scaled by size. Examples of chromosomal rearrangements are marked by roman numerals. C) Phylogenetic heatmap of microsynteny clusters. Columns represent species, arranged identically to the phylogeny in A). Numbers above columns correspond to clades in A). Rows represent microsynteny clusters, ordered by Ward’s minimum variance hierarchical clustering. Cells are colored by the number of genes present in each synteny cluster; white indicates a species has no genes in the given synteny cluster.

The conservation of cypriniform macrosynteny is striking compared to other fish clades, which exhibit substantially more chromosomal rearrangement. For example, in Siluriformes we find evidence of complex chromosomal fusions and translocations: *Clarias gariepinus* contains two chromosomal fusions that are unique among the sampled catfishes (Fig. 2B-II,III). In other cases, there may be phylogenetic signal to the macrosynteny patterns. For instance, the catfish family Ictaluridae (represented here by the genera *Ictalurus* and *Ameiurus*) appears to have undergone an ancestral chromosomal fusion (Fig. 2B-IV). Even more extreme chromosomal rearrangements occur within Characoidei; one species, *Astyanax mexicanus*, has a massive 134 Mbp chromosome with macrosynteny homology to four separate cypriniform chromosomes (Fig. 2B-I).

### Phylogenomics

Sequence-based phylogenetic approaches produced nearly identical tree topologies with strong support values at most nodes (Fig. S2), with the following exceptions. The IQTree concatenated approach (bootstrap support = 78%) and STAG (proportion of gene trees supporting node = 1.0) based on 423 single-copy orthologs both resolved *Electrophorus electricus* (Gymnotiformes) as the sister lineage of Characoidei, whereas the ASTRAL-Pro method based on 8,731 orthogroups resolved *E. electricus* as the sister lineage of Siluriformes (local posterior probability = 1.0) with an extremely short stem internode. Within Siluriformes, the IQTree concatenated approach and ASTRAL-Pro resolve *Clarias gariepinus* as the sister lineage of all other sampled catfishes, whereas STAG resolves *C. gariepinus* sister to Ictaluridae, with 0.91 proportion of input trees supporting the node.

All three sequence-based approaches resolve the same phylogenetic topology for Cypriniformes (Fig. S2, Fig. 1C). In particular, there is maximal support for *Gyrinocheilus* as the sister lineage of all other cypriniforms across all three analyses. The sequence-based approaches resolve Cobitoidei (loaches) sister to Cyprinoidei (minnows and carps), though with a very short stem internode and poor support (proportion of gene trees supporting node = 0.25) in the STAG analysis (Fig. S2, Fig. 1C).

Our initial microsynteny phylogenetic analyses produced topologies that were highly inconsistent with both our sequence-based methods and prior phylogenetic work (Fig. 1B). For instance, none of the cypriniform subclades were resolved as monophyletic (Fig. S3). The initial microsynteny tree contained four taxa with noticeably longer branch lengths: *Prochilodus magdalenae* (NCBI GCA_024036415.1), *Anabarilius grahami* (NCBI GCA_003731715.1), *Danionella translucida* (NCBI GCA_007224835.1), and *Bagarius yarrelli* (NCBI GCA_005784505.1). These four taxa all had fragmented genome assemblies (scaffold N50 < 5 Mb) and ranked lowest in BUSCO completeness (Fig. S3). Additionally, the annotations for these four assemblies were all user-submitted. Taken together, this suggests that a combination of poor assembly contiguity and low annotation quality may have confounded the microsynteny phylogenetic analyses. Therefore, we removed the four problematic taxa from the dataset for the final microsynteny phylogenetic analyses.

Microsynteny phylogenetic analyses with the filtered dataset were based on a set of 51,040 microsynteny clusters. 4,639 clusters were invariant, 46,401 clusters were variable, and 42,472 clusters were phylogenetically informative. Analyses produced a tree topology that is remarkably consistent with the sequence-based approaches (Fig. S4, Fig 1C), with a few notable exceptions. First, the microsynteny phylogeny resolved *Electrophorus electricus* (Gymnotiformes) as the sister lineage of Characoidei and Siluriformes (bootstrap support = 100%). Like the IQTree and ASTRAL-Pro sequence-based trees, microsynteny analyses also support *C. gariepinus* as sister to all other siluriforms (boostrap support = 100%). Curiously, the microsynteny phylogeny did not resolve the two sampled species of *Pangasianodon* as monophyletic. Given that *Pangasius djambal* and *Pangasianodon gigas* had user-submitted annotations (both by GenoFish) as opposed to the RefSeq annotation for *Pangasianodon hypophthalmus*, their grouping in the tree may be driven by small differences between different annotation approaches rather than true phylogenetic signal for differences in synteny. All other outgroup relationships were congruent with the sequence-based trees.

The microsynteny phylogeny strongly supports *Gyrinocheilus* as the sister lineage to all other Cypriniformes (bootstrap support = 100%), consistent with the sequence-based approaches. The monophyly of Cypriniformes is supported by 1,201 synteny synapomorphies and the monophyly of cypriniforms exclusive of *Gyrinocheilus* is supported by 79 synteny synapomorphies. However, unlike the sequence-based inference, the microsynteny analysis resolves Catostomidae (suckers) sister to Cyprinoidei (minnows and carps) (bootstrap support = 100%).

### Sequence and Synteny Topology Testing

To further examine the phylogenetic backbone of Cypriniformes, we examined several metrics of topological support. First, we calculated gene and site concordance factor values using the concatenated IQTree topology as the guide tree. Gene tree concordance factors were high for most deep nodes in the phylogeny, with two notable exceptions (Fig. S5). First, gene concordance factors for the phylogenetic resolution of Gymnotiformes, Characoidei, and Siluriformes were split between the three possible topologies: 34.5% for Gymnotiformes + Characoidei, 31.2% for Characoidei + Siluriformes, and 27.1% for Gymnotiformes + Siluriformes. Site concordance factors were also largely split: 35.6% for Gymnotiformes + Characoidei, 33.0% for Characoidei + Siluriformes, and 31.3% for Gymnotiformes + Siluriformes. However, both concordance factors marginally supported the same topological resolution of a sister relationship between Gymnotiformes and Characoidei.

The other deep topological relationship with poor concordance factor support was the resolution of Catostomidae, Cyprinoidei, and Cobitoidei within Cypriniformes. Here, gene concordance factors are evenly split between Cobitoidei + Cyprinoidei and Cobitoidei + Catostomidae, with slightly lower support for Catostomidae + Cyprinoidei (Fig. 3A). However, unlike the gene concordance factors for Gymnotiformes, Characoidei, and Siluriformes, the 27% of gene trees were uninformative about the three alternate topologies for Catostomidae, Cyprinoidei, and Cobitoidei (Fig. 3A), indicative of poor phylogenetic signal in the genes. Site concordance factors slightly favored Catostomidae + Cyprinoidei by a slight margin over the other topologies (Fig. 3A).

**Figure 3.**
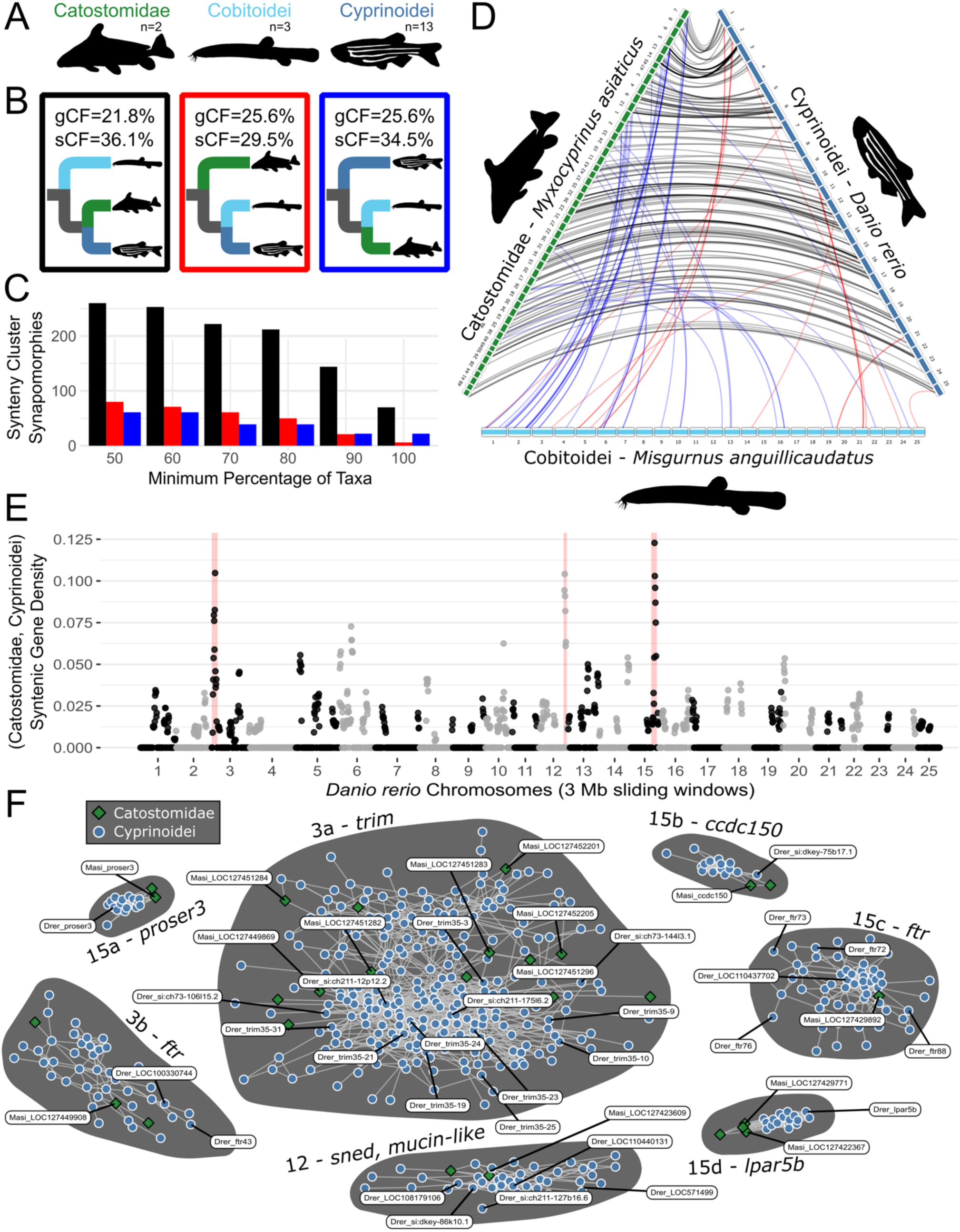
Interrogating a recalcitrant phylogenetic node using microsynteny. A) Three cypriniform subclades, number of sampled taxa indicated. B) Three alternate resolutions of Cobitoidei, Catostomidae, and Cyprinoidei. Gene concordance factor (gCF) and site concordance factor (sCF) support for each topology indicated. Boxes around each topology are colored to match bars in C), the number of microsynteny clusters supporting each topology across a range of taxon filtering thresholds. D) Clade-specific microsynteny clusters shared by at least 90% of the ingroup taxa. Each link represents a syntenic gene pair between the representative genomes for each clade. Links are colored by the topology they support (panel B). E) *Danio rerio* Manhattan plot of the normalized microsynteny gene density (3 Mb sliding windows, 0.5 Mb slide size) supporting the sister relationship between Catostomidae and Cyprinoidei. Red highlights regions on chromosomes 3, 12, and 15 with concentrations of microsynteny synapomorphies. F) These regions are comprised of 2, 1, and 4 microsynteny clusters, respectively. Each point indicates a gene from a specific species, blue circles are cyprinoids, green diamonds are catostomids. Links indicated pairwise synteny. Clusters are annotated with the primary representative gene or gene family name.

We used the microsynteny phylogenetic signal to further examine topological support for the resolution of Catostomidae, Cyprinoidei, and Cobitoidei. For each of the three rooted topologies, we counted the number of synteny synapomorphies (synteny clusters exclusive to the focal clade). Across a range of minimum taxa thresholds, the clade of Catostomidae + Cyprinoidei had between three to 12 times more synteny synapomorphies than the alternate topologies (Fig. 3C). When we compared representative taxa for each clade, we found that the synteny synapomorphies for Catostomidae + Cyprinoidei are present on nearly every chromosome (Fig. 3D,E). By contrast, Cobitoidei + Cyprinoidei and Cobitoidei + Catostomidae synteny synapomorphies are absent on many chromosomes (Fig. 3D).

### Genomic Hotspots of Microsynteny Signal

To determine whether microsynteny phylogenetic signal for the sister relationship between Catostomidae and Cyprinoidei was concentrated in particular genomic regions or dispersed broadly across the genome, we used a sliding window approach to calculate the proportion of syntenic genes along the *Danio rerio* reference chromosomes (normalized by the total number of genes in each window). While there were synteny synapomorphies on every chromosome for the representative species (Fig. 3D), three regions on *D. rerio* chromosomes 3, 12, and 15 stood out (Fig. 3E). The outlier region on chromosome 3 contains two synteny clusters unique to the clade comprising Catostomidae and Cyprinoidei. One of these clusters is comprised of 46 genes in the *ftr* family, while the other more substantial cluster is comprised of 274 genes in the *trim* family (Fig. 3F, Table S3). The chromosome *Danio rerio* 12 outlier region contains a single synteny cluster comprised of 44 genes, consisting mostly of *sned1* and *mucin-like* genes (Fig. 3F, Table S3). Lastly, the outlier region of chromosome 15 contains four synteny clusters: three small clusters comprised exclusively of the genes *proser3, ccdc150,* and *lpar5b*, as well as another cluster containing 59 genes in the *ftr* family (Fig. 3F, Table S3). To summarize, all three of the outlier regions (on chromosomes 3, 12, and 15) appear to contain tandemly-duplicated gene family expansions as mechanistic drivers of synteny synapomorphies.

In addition to the clade comprising Catostomidae and Cyprinoidei, we analyzed clade-specific synteny cluster synapomorphies across five clades of varying evolutionary ages: Ostariophysi, Cypriniformes, Cyprinoidei, Catostomidae, and Cobitoidea (Fig. 4). The number of these synteny synapomorphies varied considerably among the clades, with Catostomidae having the most (1,209 clusters) and Cyprinoidei having the fewest (133 clusters). The distribution of these synapomorphies also showed variable patterns. Clades like Ostariophysi and Cypriniformes exhibited a generally high background of about 1,000 clusters, but with no distinguishable hotspots. In contrast, Cyprinoidei and Cobitoidea had low background levels of only a few hundred clusters, although Cyprinoidei was distinguished by a single, strong outlier peak of high synteny cluster density.

**Figure 4.**
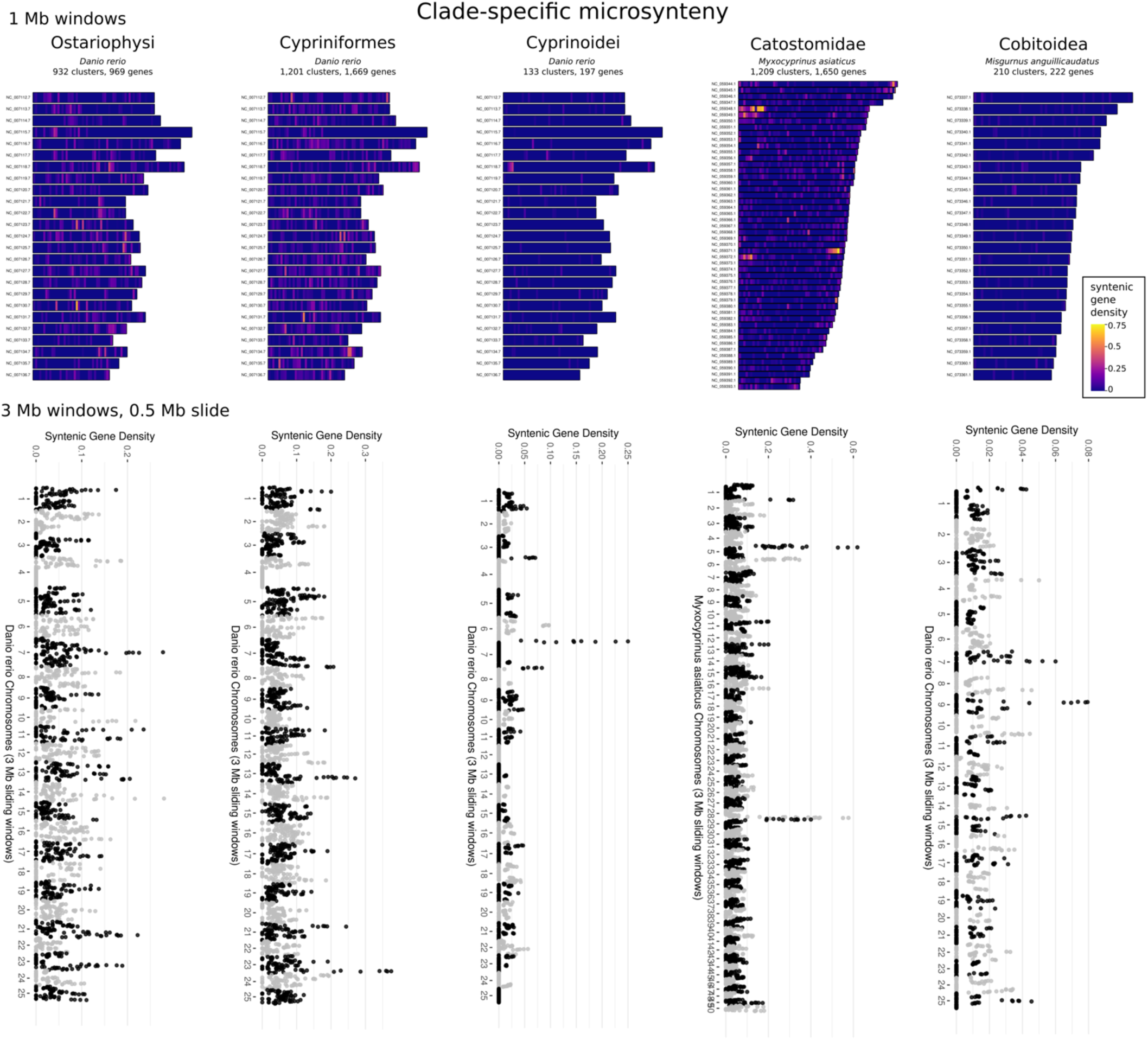
A) (top row) Chromosomal heatmaps of clade-specific normalized syntenic gene density in 1 Mb windows. Heatmap scale is consistent across all clades. Number of clade-specific microsynteny clusters (and the genes comprising those clusters) is noted. Genomes shown are Danio rerio (Ostariophysi, Cypriniformes, and Cyprinoidei), *Myxocyprinus asiaticus* (Catostomidae), and *Misgurnus anguillicaudatus* (Cobitoidea). B) (bottom row) Manhattan plots showing 3 Mb sliding windows (0.5 Mb slide size) of clade-specific normalized syntenic gene density.

Catostomidae presented both a high background number of clusters and several prominent outlier peaks. Genes within these hotspots are concentrated on four chromosomes that represent two pairs of homeologs from the catostomid-specific whole genome duplication event (“Cat-4R”) (Krabbenhoft et al. 2021). Each pair of these homeologous chromosomes contains microsynteny clusters with genes associated with testes and ovarian development, including tandemly duplicated blocks of zinc-finger genes. This pattern suggests that lineage-specific conservation of sex-associated chromosomal regions may be a novel source of microsyntenic phylogenetic signal. This finding is significant given that cypriniform fishes are known for high rates of turnover in their sex chromosomes and sex-determining regions and typically lack heteromorphic sex chromosomes (Wilson et al. 2014; Caeiro-Dias et al. 2023; Schlebusch et al. 2025). This turnover could contribute to the phylogenetic signal and underscores the importance of considering the sex of individuals used for reference genomes in microsynteny analyses.

## DISCUSSION

### A Phylogenomic Framework for Cypriniformes

The evolutionary history of Cypriniformes, one of the most species-rich orders of freshwater fishes, has been characterized by persistent uncertainty, particularly at its deepest nodes (Conway et al. 2010; Stout et al. 2016). This phylogenetic ambiguity has hindered the development of a stable classification and impeded progress in understanding the macroevolutionary processes that generated the order’s immense diversity. Resolving deeper evolutionary relationships of Cypriniformes also has important implications for biomedical research, as the order contains the Zebrafish, *Danio rerio,* the most well-studied fish model organism (Meyers 2018). Here we provide the first genome assembly for the family Gyrinocheilidae and use both sequence-based and microsynteny-based phylogenomic approaches to clarify the backbone of the cypriniform tree of life. Our results resolve long-standing topological questions and uncover a novel, unexpected relationship among major cypriniform lineages, fundamentally reshaping our understanding of the evolutionary history of this group’s early diversification.

Gyrinocheilidae is perhaps the most unusual of the four major cypriniform lineages. It is monogeneric and species-poor, containing only three recognized species. We also found that *G. aymonieri* has the smallest genome of any sampled cypriniform, about 200 Mb shorter than the next closest species, along with low repeat content in the genome and chromosomal fusion versus outgroup taxa. This lineage is also unusual from a phylogenetic and taxonomic perspective. Early taxonomists included Gyrinocheilidae in the superfamily Cobitoidea along with loaches and suckers (reviewed in Conway et al. 2010), but more recently Cobitoidea has been reclassified to only contain the loach families (as Cobitoidei, Kottelat 2012; Stout et al. 2016). This is due to the lack of phylogenetic clarity regarding loaches, suckers, and Gyrinocheilidae. Gyrinocheilidae has been variously resolved as the sister lineage of Catostomidae or the sister lineage of loaches (Fig. 1B). Most recently, molecular analyses by Stout & Tan et al. (2016) produced conflicting phylogenetic placements of Gyrinocheilidae (Fig. 1B). Given the criticisms of species tree methods (Gatesy and Springer 2014; Tonini et al. 2015), the authors prioritized their concatenated analysis, which identified Gyrinocheilidae as the sister lineage to all other cypriniforms.

Both our sequence-based and synteny-based analyses of whole genomes consistently infer Gyrinocheilidae to occupy the pivotal phylogenetic position as the outgroup to all other cypriniforms (Fig. 1C, Fig. S2, Fig. S4). Concordance factors also support this topology, with 58.5% of gene trees and 40.7% of alignment sites resolving a cypriniform subclade exclusive of Gyrinocheilidae (Fig. S5). Congruent phylogenetic signal from thousands of protein-coding gene sequences and the genomic arrangement of those genes provides two distinct lines of evidence supporting this phylogenetic hypothesis. This result is supported by at least one morphological synapomorphy: in all cypriniforms except Gyrinocheilidae, the pharyngeal teeth on the fifth ceratobranchial element are ankylosed to the bone while Gyrinocheilidae completely lacks pharyngeal teeth (Fink and Fink 1981). Previous authors considered this a synapomorphy of cypriniforms that was secondarily lost in Gyrinocheilidae (Conway et al. 2010). However, the placement of Gyrinocheilidae sister to all other cypriniforms provides a more parsimonious explanation for the evolution of this trait.

While the placement of Gyrinocheilidae confirms a previously hypothesized relationship, our analyses reveal a novel and unexpected topology for the remaining cypriniform lineages. In contrast to the results from our sequence-based methods and all prior molecular studies, our microsynteny analysis provides strong support for a sister relationship between Catostomidae and Cyprinoidei. This conclusion is supported by a striking disparity in the number and genomic distribution of phylogenetically informative microsynteny clusters. Across a range of filtering thresholds, the number of shared, derived synteny clusters (synteny synapomorphies) supporting Catostomidae + Cyprinoidei is three to twelve times greater than for the alternative topologies (Fig. 3C, D). The microsynteny phylogeny directly contradicts our own sequence-based phylogenetic analyses, which inferred a weakly supported sister relationship between Cobitoidei and Cyprinoidei (Fig. 1D, Fig. S2), consistent with concatenated analysis by Stout & Tan et al. (2016). The weak support for this relationship is driven by lack of phylogenetic signal in the underlying data (Fig. 3B), a classic characteristic of short phylogenetic internodes produced by rapid lineage splitting.

Our revised phylogenetic backbone for Cypriniformes has important implications, particularly for biomedical research that relies on the Zebrafish, *Danio rerio*, a member of Cyprinoidei. Our results reposition the suckers (Catostomidae) as a closer outgroup to the minnows and carps (Cyprinoidei) than the loaches (Cobitoidei). This suggests that a catostomid species, rather than a cobitid, would be a more phylogenetically appropriate comparative model for investigating the evolutionary origins of traits specific to the Cyprinoidei lineage, including developmental, genomic, and physiological features relevant to the Zebrafish model system. Of course, catostomids have undergone their own recent genome duplication (Krabbenhoft et al. 2021), which could complicate comparative evolutionary approaches.

The difficulty in resolving the cypriniform backbone is mirrored at a deeper evolutionary scale in the relationships among the major lineages of the superorder Ostariophysi. Specifically, the Siluriformes (catfishes), Characoidei and Citharinoidei (characins), and Gymnotiformes (knifefishes) are a classic example of a contentious phylogenetic problem in the fish tree of life (reviewed in Dornburg and Near 2021). Even large molecular datasets have failed to produce a consensus topology (Arcila et al. 2017; Chakrabarty et al. 2017; Melo et al. 2022). Unsurprisingly, our sequence-based analyses are also inconclusive (Fig. S2) and concordance factors show split support (Fig. S5). However, our microsynteny topology (Fig. 2A, S4) is consistent with (what some consider) the most well-supported molecular hypotheses (Dornburg and Near 2021). Unfortunately, the lack of available genomes for Citharinoidei prevents us from developing a more complete phylogenomic answer. Future efforts should develop high-quality genome assemblies for Citharinoidei and examine whether microsynteny holds key phylogenetic signal to resolve these relationships.

### The Phylogenetic Power of Microsynteny

The conflicting results between our sequence and synteny analyses for the phylogenetic relationship between Catostomidae, Cobitoidei, and Cyprinoidei provide a powerful case study for evaluating the relative strengths and weaknesses of different approaches to phylogenomics. From a macrosynteny (chromosome-level) perspective, there appears to be very little phylogenetic signal for Cypriniformes. Despite ∼100 million years of evolution (Near et al. 2012; Hughes et al. 2018) that includes several independent origins of polyploidy (Li and Guo 2020; Krabbenhoft et al. 2021; Lv et al. 2025), most available cypriniform genomes exhibit a 1:1 (or 2:1) orthologous relationship among chromosomes with few major rearrangements (Fig. 2B). However, our findings demonstrate that microsynteny can be a useful tool for resolving difficult nodes while also highlighting the need for high-quality reference genomes, offering lessons for the design of future phylogenomic studies.

The short internodes separating the major cypriniform clades in our sequence-based trees combined with the split conflicting concordance factor support may indicate a history of rapid diversification. Ancient rapid radiations are a well-known challenge for phylogenetic methods (Whitfield and Lockhart 2007), since the number of substitutions retained is related to internode branch lengths and time since divergence (Townsend 2007) and the degree of incomplete lineage sorting (ILS) producing gene tree conflict is inversely proportional to internode branch lengths (Degnan and Rosenberg 2009). We suggest that microsynteny may provide cleaner phylogenetic signal in such cases. Compared to point mutations, gene collinearity is the product of more complex evolutionary events such as translocations, inversions, and gene duplications. Modification of synteny is considered a “rare genomic change” that generally occurs at a slower rate than point mutations (Rokas and Holland 2000; Steenwyk and King 2024b).Yet genomic structural variation can also segregate quickly in a population, potentially limiting the effects of ILS (Rokas and Holland 2000). However, we note that structural genomic variation (and associated changes in synteny) can also be maintained by selection as stable polymorphisms (e.g. Sodeland et al. 2016; Kratochwil et al. 2022), which could produce misleading phylogenetic signal. Homoplasy (and resulting long-branch attraction) is also much less likely to occur in microsynteny datasets, as convergent evolution of collinear genes is less likely given the large number (tens of thousands) of potential character states. Consequently, a shared syntenic block is more probable to be a true, derived character (a synapomorphy) uniting a clade, thereby providing a more robust signal that cuts through the noise of ILS. The difference in support for Catostomidae + Cyprinoidei topology between our sequence and synteny analyses serves as a powerful empirical demonstration of this principle. It is unclear if any morphological synapomorphies support this novel phylogenetic topology, as morphological trees generally have low support for deep cypriniform relationships (Stout et al. 2016).

While microsynteny analyses have power the to resolve difficult phylogenetic problems like the cypriniform backbone, our study also delivers a cautionary tale. Our initial microsynteny analysis of 43 genomes from NCBI produced a phylogeny in which two of the major cypriniform subgroups, Cyprinoidei and Cobitoidei, were resolved as polyphyletic (Fig. S3A). In contrast, sequence-based phylogenetic analyses of the same 43 genomes resolved the four major Cypriniform subclades (Fig. S2), consistent with all prior molecular analyses (Fig. 1B). Closer examination revealed that the phylogenetic discordance in the microsynteny analysis was driven by the inclusion of just four taxa whose genomes and gene models were of lower quality (Fig. S3B-D), characterized by highly fragmented assemblies (scaffold N50 < 5 Mb) and incomplete gene annotations (<90% complete BUSCO scores).

This outcome highlights a limitation of synteny-based phylogenetic approaches. These methods are more sensitive to the quality of the input genomic data than sequence-based inference. A fragmented assembly artificially breaks up regions of true gene collinearity, creating false signals of rearrangement (Liu et al. 2018) that can be misinterpreted as phylogenetic signal. Similarly, incomplete or inaccurate gene annotations increase the difficulty of identifying true orthologs, leading to an inaccurate synteny network. For decades, phylogenetic studies have typically prioritized increasing taxon sampling to improve accuracy, a strategy well-suited to sequence-based methods that are relatively robust to the inclusion of draft-quality genomes (e.g. Zhang et al. 2019). Our work demonstrates that for microsynteny analyses, adding more taxa may not always be advisable. The inclusion of even a small fraction (in our case, <10%) of low-quality genomes confounded phylogenetic inference. This implies that for resolving deep, recalcitrant nodes with microsynteny, generating a smaller number of high-quality, chromosome-level genomes for key taxa may be more effective than generating draft-quality genomes for a broader array of taxa.

### Distribution of Phylogenetically Informative Synteny

Phylogenetic signal from microsynteny synapomorphies is not uniformly distributed across the genome. While phylogenetically informative microsynteny clusters are generally widespread and found on most chromosomes in the representative species, some clades exhibit distinctive genomic hotspots of phylogenetic signal. For the clade comprised of Catostomidae and Cyprinoidei, the three largest hotspots contain both large and small synteny clusters (Fig. 3D-F). The large synteny clusters are comprised of many copies from the same gene family, perhaps indicating localized, clade-specific tandem duplication events (Fig. 3F, Table S3). In contrast, some smaller synteny clusters contain only one or two copies of each gene, possibly signaling clade-specific gene loss or translocation (Fig. 3F, Table S3). Other clades also exhibit such hotspots, most notably Catostomidae, where two homeologous chromosome pairs show elevated microsynteny phylogenetic signal (Fig. 4). The microsynteny method we employed cannot distinguish the underlying structural genomic changes that produced phylogenetic patterns. Additionally, it is unclear whether these hotspots of synteny phylogenetic signals simply represent hotspots of genomic structural change (Lin and Gokcumen 2019). Regardless, changes in synteny driven by changes in genome structure can have dramatic evolutionary consequences, as a single SV impacts far more base pairs and genes than a single point mutation (Kloosterman et al. 2015; Mirarab et al. 2024). Thus, the evolutionary drivers of these microsynteny patterns and their broader implications remains a tantalizing avenue for future research. These results extend beyond phylogenomics, offering both preliminary insights into the evolutionary success of the hyperdiverse cypriniform fishes and a starting point for future functional genomics studies

## ACKNOWLEDGEMTS

We would like to thank C. A. Osborne, K. Louisor, L. N. Gray, and V. A. Albert for valuable feedback on earlier versions of this work.

## FUNDING

Support for this work was provided by laboratory startup funds from the University at Buffalo to TJK. DJM was supported in part by an NSF PRFB (award #2109761).

## DATA AVAILABILITY

The *Gyrinocheilus aymonieri* genome assembly (NCBI PRJNA784511) will be released on NCBI upon publication. Nanopore and Illumina DNA sequence data and the *G. aymonieri* genome assembly are available at NCBI PRJNA763389 (SRR15902715 - Nanopore reads, SRR15902716 - Illumina reads). The genome annotation for *G. aymonieri* will be made available on Dryad upon publication. Genome assembly, genome annotation, and analysis scripts are available on GitHub https://github.com/KrabbenhoftLab/Gyrinocheilus_genome.

---

**Figure S1.**
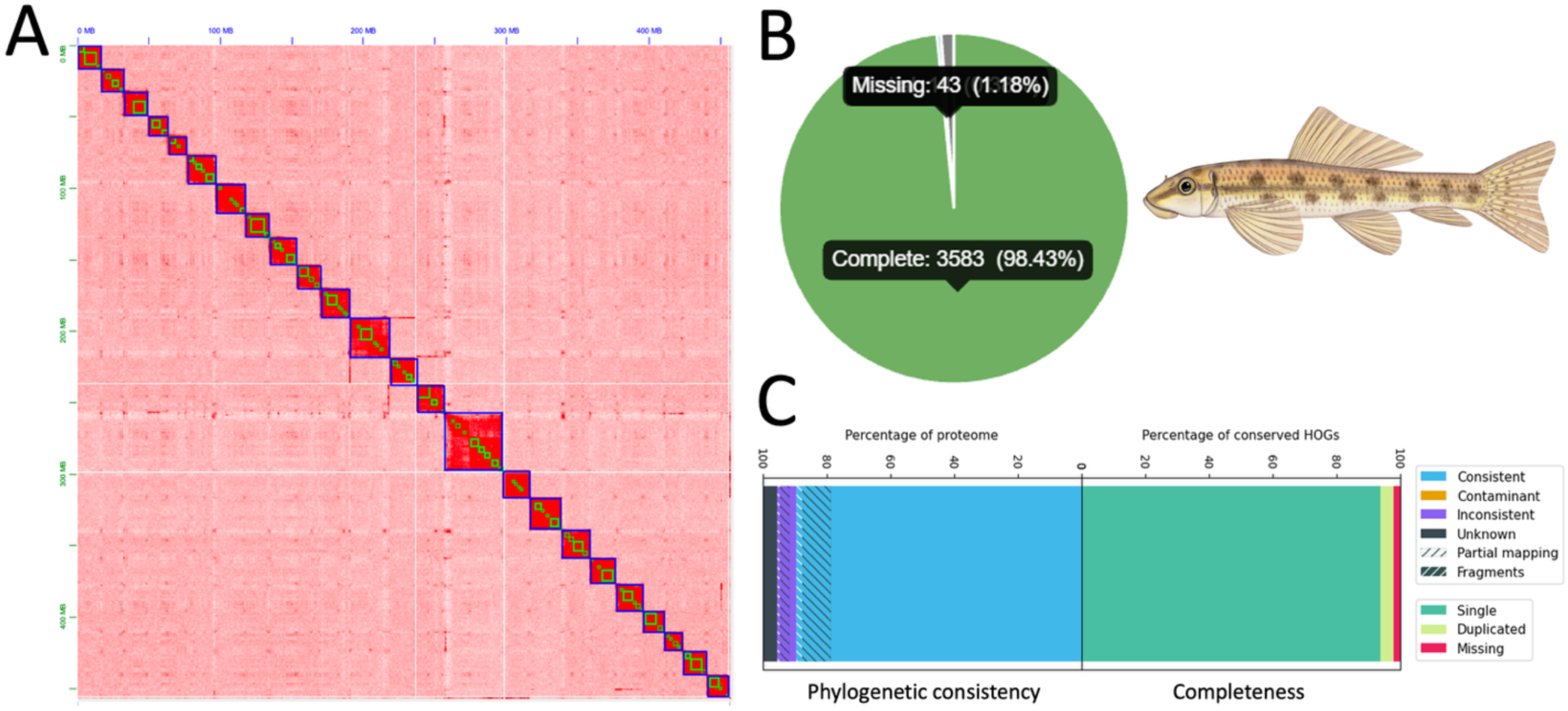
The *Gyrinocheilus aymonieri* genome. A) Hi-C heatmap of the assembled scaffolds. Green boxes indicate contig boundaries, blue boxes indicate scaffold boundaries. B) BUSCO pie chart and C) OMArk results for the predicted *G. aymonieri* genes. Artwork provided by Joseph R. Tomelleri.

**Figure S2.**
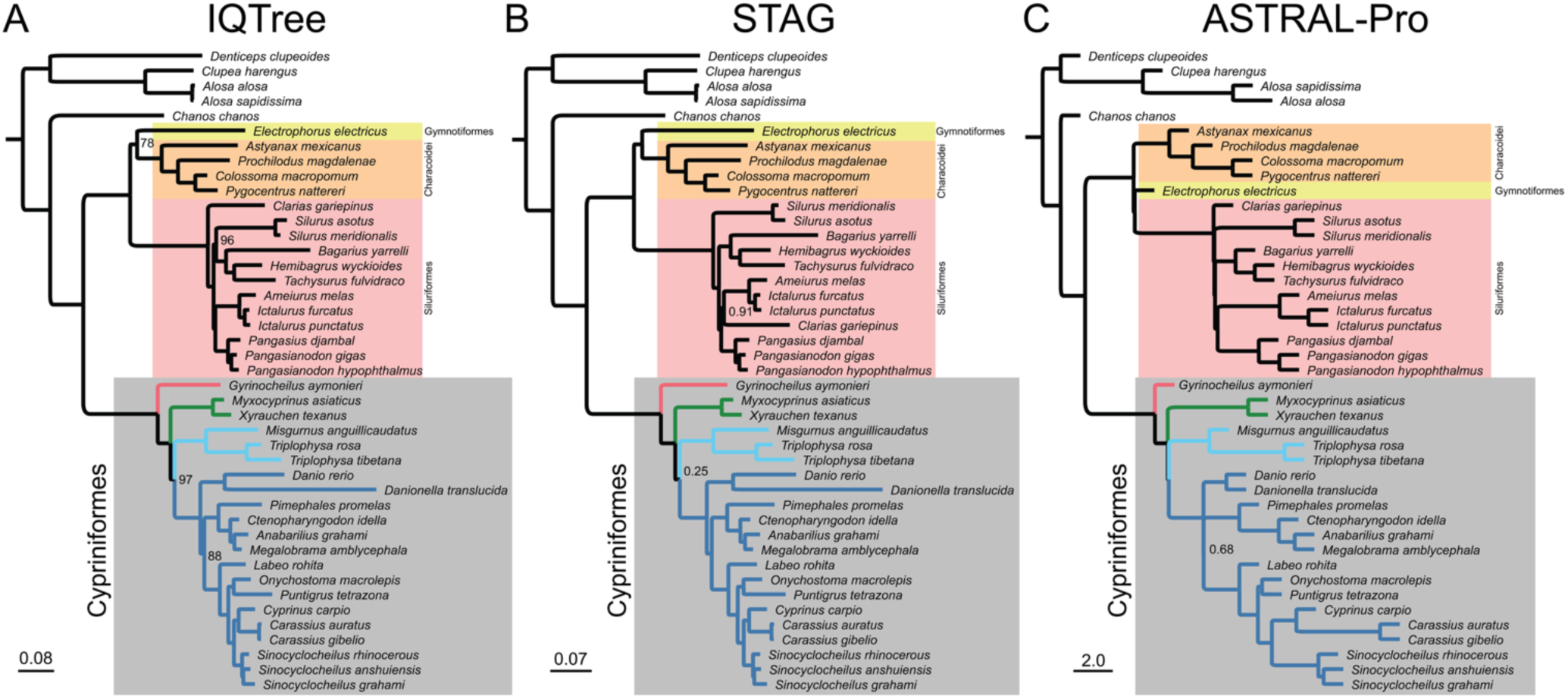
Sequence based phylogenies from A) a concatenated maximum likelihood approach with IQTree (with ultrafast bootstrap support values), B) a consensus species tree approach with STAG (with nodal proportion of gene trees support) and C) a coalescent-based species tree approach with ASTRAL-Pro (with local posterior probabilities). Branches within Cypriniformes are colored following Fig. 1. Nodes with less than maximal support values are labeled. Branches lengths are in unites of expected number of substitutions per site (IQTree and STAG) or coalescent units (ASTRAL-Pro). Note that ASTRAL does not estimate terminal branch lengths.

**Figure S3.**
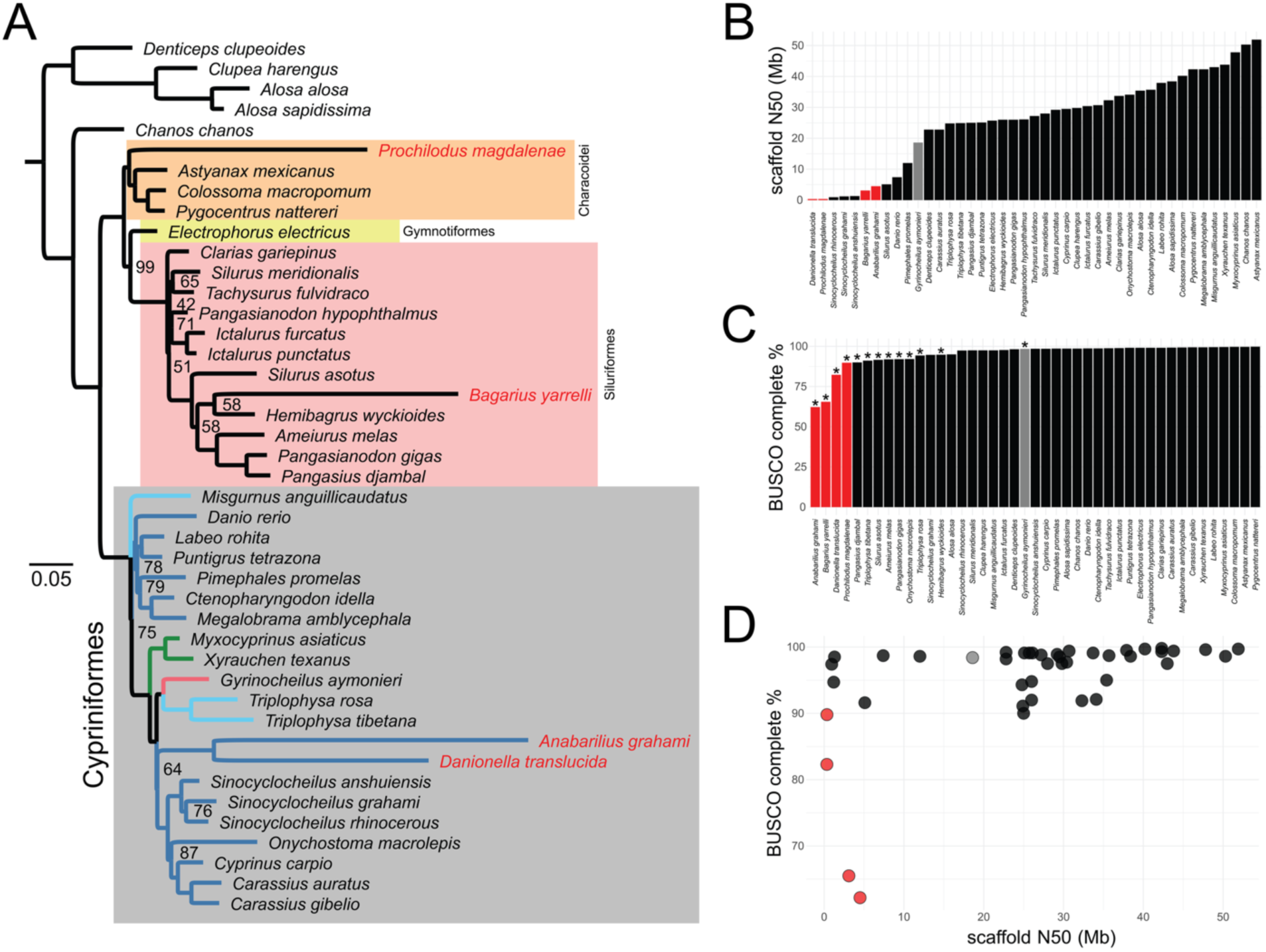
Low-quality genome assemblies and annotations result in spurious phylogenetic relationships. A) Microsynteny tree topology using all genomes included in the sequence-based phylogenetic analyses. Four species with long terminal branches are highlighted in red. Cypriniform branches are colored by clade (Fig. 1). B) Species ranked by genome contiguity (scaffold N50 [Mbp]) and C) species ranked by annotation quality (BUSCO % complete). Asterisks indicate user-submitted annotations, all other annotations produced by NCBI. Red bars in B) and C) correspond to the four long-branch taxa in A). D) Scatter plot of genome annotation quality versus assembly contiguity. Red points correspond to the four long-branch taxa in A). Our *Gyrinocheilus aymonieri* assembly and annotation is highlighted in gray in B-D).

**Figure S4.**
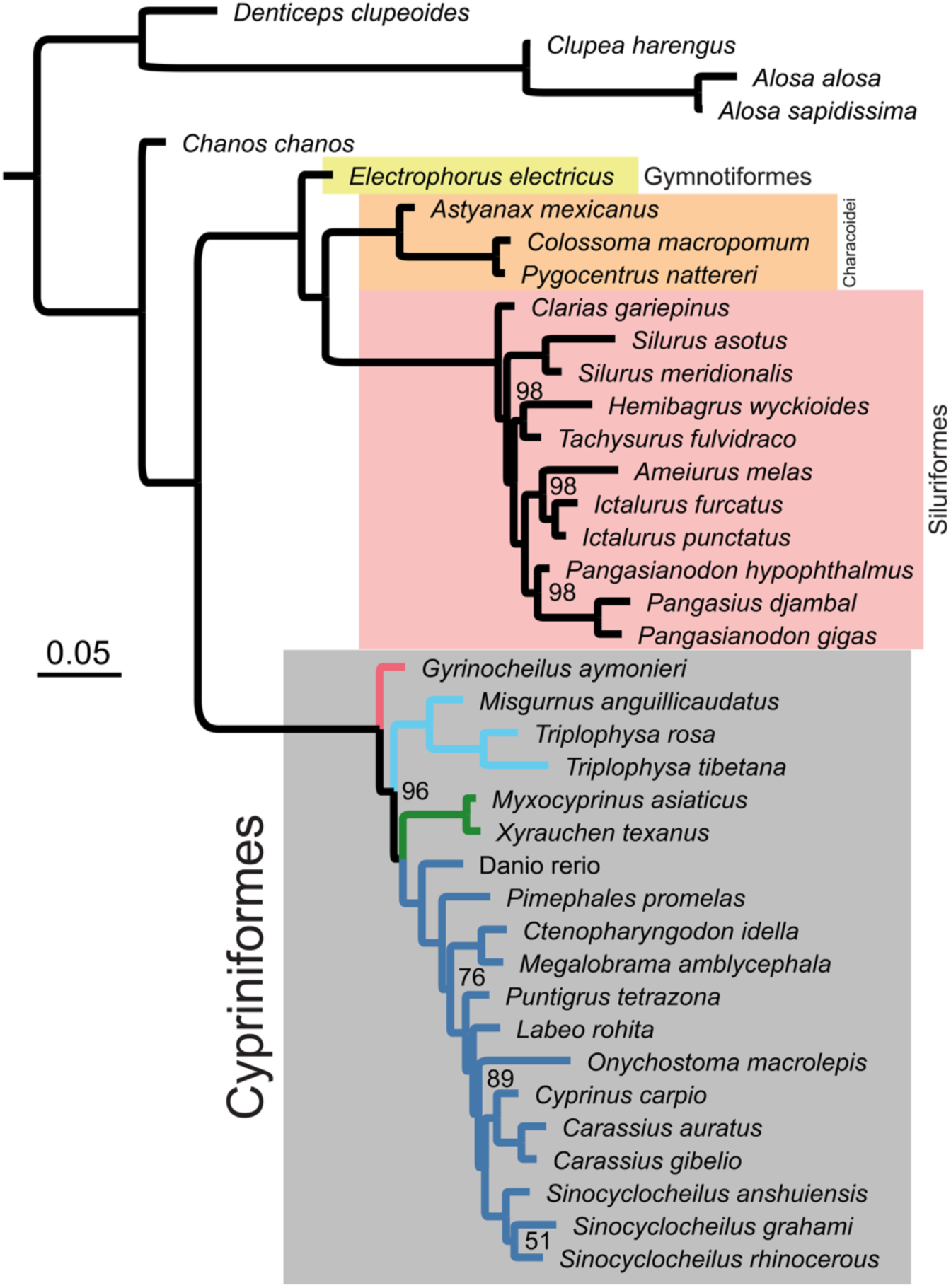
Microsynteny phylogeny for Otocephala. Branch lengths are proportional to the expected number of changes per character (synteny cluster). Nodes with bootstrap support values <100% are labeled. Cypriniform branches are colored by clade (Fig. 1).

**Figure S5.**
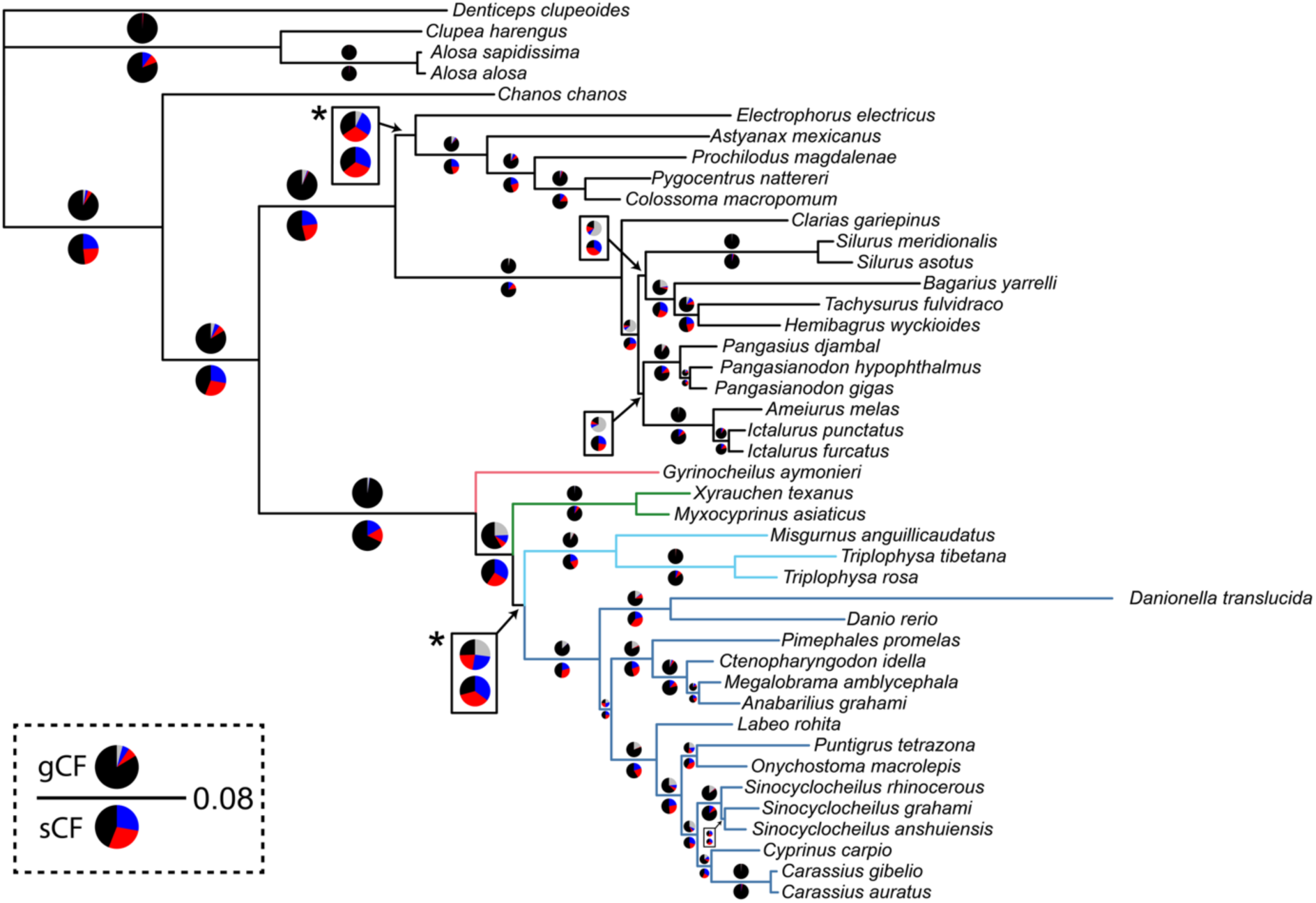
Concordance factors (CF) values visualized on the ASTRAL-Pro tree topology. Pie charts above branches represent gene CFs (gCFs), while pie charts below represent site CFs (sCFs). Black indicates agreement with the displayed tree, red and blue indicate disagreement, and gray indicates no support or noise. CF pie charts are scaled for visual clarity. Some CF pie charts are offset from the associated branch due to space limitations. Asterisks indicate two deep nodes with indecisive CFs discussed in the text. Branch lengths are in coalescent units, branch length scale indicated in dashed box. Cypriniform branches are colored by clade (Fig. 1).

**Table S1.** NCBI genome summary stats. See TableS1.csv.

**Table S2.** Number of retained orthogroups (single-copy for diploid taxa, maximum of two copies for polyploid taxa) at different missing taxa thresholds.

| Minimum<br># taxa | Maximum %<br>missing taxa | # of retained<br>orthogroups |
| --- | --- | --- |
| 34 | 21 | 11,700 |
| 35 | 19 | 11,428 |
| 36 | 16 | 11,064 |
| 37 | 14 | 10,533 |
| 38 | 12 | 9,817 |
| 39 | 9 | 8,713 |
| 40 | 7 | 7,139 |
| 41 | 5 | 5,128 |
| 42 | 2 | 2,777 |
| 43 | 0 | 831 |

**Table S3.** List of genes for each of the synteny clusters in the Catostomidae + Cyprinoidei microsynteny hotspots in Fig. 3E,F. See TableS3.xlsx.

## Notes

### Competing Interest Statement

The authors have declared no competing interest.

## REFERENCES

1. Almeida-Silva F., Zhao T., Ullrich K.K., Schranz M.E., Van de Peer Y. 2023. syntenet: an R/Bioconductor package for the inference and analysis of synteny networks. Bioinformatics. 39:btac806.

2. Arcila D., Ortí G., Vari R., Armbruster J.W., Stiassny M.L.J., Ko K.D., Sabaj M.H., Lundberg J., Revell L.J., Betancur-R R. 2017. Genome-wide interrogation advances resolution of recalcitrant groups in the tree of life. Nat. Ecol. Evol. 1:20.

3. Armstrong J., Hickey G., Diekhans M., Fiddes I.T., Novak A.M., Deran A., Fang Q., Xie D., Feng S., Stiller J., Genereux D., Johnson J., Marinescu V.D., Alföldi J., Harris R.S., Lindblad-Toh K., Haussler D., Karlsson E., Jarvis E.D., Zhang G., Paten B. 2020. Progressive Cactus is a multiple-genome aligner for the thousand-genome era. Nature. 587:246–251.

4. Britz R., Conway K.W. 2011. The Cypriniformes Tree of Confusion. Zootaxa. 2946:73–78.

5. Brůna T., Lomsadze A., Borodovsky M. 2024. GeneMark-ETP significantly improves the accuracy of automatic annotation of large eukaryotic genomes. Genome Res. 34:757–768.

6. Buchfink B., Reuter K., Drost H.-G. 2021. Sensitive protein alignments at tree-of-life scale using DIAMOND. Nat. Methods. 18:366–368.

7. Caeiro-Dias G., Osborne M.J., Waterman H.M., Krabbenhoft T.J., Turner T.F. 2023. Limited evidence for extensive genetic differentiation between X and Y chromosomes in Hybognathus amarus (Cypriniformes: Leuciscidae). J. Hered. 114:470–487.

8. Chakrabarty P., Faircloth B.C., Alda F., Ludt W.B., Mcmahan C.D., Near T.J., Dornburg A., Albert J.S., Arroyave J., Stiassny M.L.J., Sorenson L., Alfaro M.E. 2017. Phylogenomic Systematics of Ostariophysan Fishes: Ultraconserved Elements Support the Surprising Non-Monophyly of Characiformes. Syst. Biol. 66:881–895.

9. Conway K.W. 2011. Osteology of the South Asian Genus Psilorhynchus McClelland, 1839 (Teleostei: Ostariophysi: Psilorhynchidae), with investigation of its phylogenetic relationships within the order Cypriniformes. Zool. J. Linn. Soc. 163:50–154.

10. Conway K.W., Hirt M.V., Yang L., Mayden R.L., Simons A.M. 2010. Cypriniformes: Systematics & Paleontology: Festschrift in honor of G. Arratia. In: Schultze H.P., Nelson J.S., Wilson M.V.H., editors. Origin and Phylogenetic Interrelationships of Teleosts. p. 295–316.

11. Dainat J., Cannoodt R., Soares A., Ruano D.G., Hereñú D., Murray Dr.K.D., Davis E., Ugrin I., Crouch K., Soler L., Agostinho N., pascal-git, Zollman Z., tayyrov. 2025. NBISweden/AGAT: AGAT v1.5.1.

12. Degnan J.H., Rosenberg N.A. 2009. Gene tree discordance, phylogenetic inference and the multispecies coalescent. Trends Ecol. Evol. 24:332–340.

13. Ding Y.-M., Pang X.-X., Cao Y., Zhang W.-P., Renner S.S., Zhang D.-Y., Bai W.-N. 2023. Genome structure-based Juglandaceae phylogenies contradict alignment-based phylogenies and substitution rates vary with DNA repair genes. Nat. Commun. 14:617.

14. Dobzhansky T., Sturtevant A.H. 1938. INVERSIONS IN THE CHROMOSOMES OF DROSOPHILA PSEUDOOBSCURA. Genetics. 23:28–64.

15. Dornburg A., Near T.J. 2021. The Emerging Phylogenetic Perspective on the Evolution of Actinopterygian Fishes. Annu. Rev. Ecol. Evol. Syst. 52:427–452.

16. Dudchenko O., Batra S.S., Omer A.D., Nyquist S.K., Hoeger M., Durand N.C., Shamim M.S., Machol I., Lander E.S., Aiden A.P., Aiden E.L. 2017. De novo assembly of the Aedes aegypti genome using Hi-C yields chromosome-length scaffolds. Science. 356:92–95.

17. Durand N.C., Shamim M.S., Machol I., Rao S.S.P., Huntley M.H., Lander E.S., Aiden E.L. 2016. Juicer Provides a One-Click System for Analyzing Loop-Resolution Hi-C Experiments. Cell Syst. 3:95–98.

18. Ellinghaus D., Kurtz S., Willhoeft U. 2008. LTRharvest, an efficient and flexible software for de novo detection of LTR retrotransposons. BMC Bioinformatics. 9:18.

19. Emms D.M., Kelly S. 2018. STAG: Species Tree Inference from All Genes. bioRxiv.:267914.

20. Emms D.M., Kelly S. 2019. OrthoFinder: phylogenetic orthology inference for comparative genomics. Genome Biol. 20:238.

21. FAO. 2024. The State of World Fisheries and Aquaculture 2024. FAO;

22. Fink S.V., Fink W.L. 1981. Interrelationships of the ostariophysan fishes (Teleostei). Zool. J. Linn. Soc. 72:297–353.

23. Flynn J.M., Hubley R., Goubert C., Rosen J., Clark A.G., Feschotte C., Smit A.F. 2020. RepeatModeler2 for automated genomic discovery of transposable element families. Proc. Natl. Acad. Sci. U. S. A.

24. Gabriel L., Brůna T., Hoff K.J., Ebel M., Lomsadze A., Borodovsky M., Stanke M. 2024. BRAKER3: Fully automated genome annotation using RNA-seq and protein evidence with GeneMark-ETP, AUGUSTUS, and TSEBRA. Genome Res. 34:769–777.

25. Gabriel L., Hoff K.J., Brůna T., Borodovsky M., Stanke M. 2021. TSEBRA: transcript selector for BRAKER. BMC Bioinformatics. 22:566.

26. Gatesy J., Springer M.S. 2014. Phylogenetic analysis at deep timescales: Unreliable gene trees, bypassed hidden support, and the coalescence/concatalescence conundrum. Mol. Phylogenet. Evol. 80:231–266.

27. Haas B.J., Salzberg S.L., Zhu W., Pertea M., Allen J.E., Orvis J., White O., Buell C.R., Wortman J.R. 2008. Automated eukaryotic gene structure annotation using EVidenceModeler and the Program to Assemble Spliced Alignments. Genome Biol. 9:R7.

28. Hao Z., Lv D., Ge Y., Shi J., Weijers D., Yu G., Chen J. 2020. RIdeogram: drawing SVG graphics to visualize and map genome-wide data on the idiograms. PeerJ Comput. Sci. 6:e251.

29. Hirt M.V., Arratia G., Chen W.-J., Mayden R.L., Tang K.L., Wood R.M., Simons A.M. 2017. Effects of gene choice, base composition and rate heterogeneity on inference and estimates of divergence times in cypriniform fishes. Biol. J. Linn. Soc. 121:319–339.

30. Hoang D.T., Chernomor O., Von Haeseler A., Minh B.Q., Vinh L.S. 2018. UFBoot2: Improving the ultrafast bootstrap approximation. Mol. Biol. Evol.

31. Howe K., Clark M.D., Torroja C.F., Torrance J., Berthelot C., Muffato M., Collins J.E., Humphray S., McLaren K., Matthews L., McLaren S., Sealy I., Caccamo M., Churcher C., Scott C., Barrett J.C., Koch R., Rauch G.-J., White S., Chow W., Kilian B., Quintais L.T., Guerra-Assunção J.A., Zhou Y., Gu Y., Yen J., Vogel J.-H., Eyre T., Redmond S., Banerjee R., Chi J., Fu B., Langley E., Maguire S.F., Laird G.K., Lloyd D., Kenyon E., Donaldson S., Sehra H., Almeida-King J., Loveland J., Trevanion S., Jones M., Quail M., Willey D., Hunt A., Burton J., Sims S., McLay K., Plumb B., Davis J., Clee C., Oliver K., Clark R., Riddle C., Elliott D., Threadgold G., Harden G., Ware D., Begum S., Mortimore B., Kerry G., Heath P., Phillimore B., Tracey A., Corby N., Dunn M., Johnson C., Wood J., Clark S., Pelan S., Griffiths G., Smith M., Glithero R., Howden P., Barker N., Lloyd C., Stevens C., Harley J., Holt K., Panagiotidis G., Lovell J., Beasley H., Henderson C., Gordon D., Auger K., Wright D., Collins J., Raisen C., Dyer L., Leung K., Robertson L., Ambridge K., Leongamornlert D., McGuire S., Gilderthorp R., Griffiths C., Manthravadi D., Nichol S., Barker G., Whitehead S., Kay M., Brown J., Murnane C., Gray E., Humphries M., Sycamore N., Barker D., Saunders D., Wallis J., Babbage A., Hammond S., Mashreghi-Mohammadi M., Barr L., Martin S., Wray P., Ellington A., Matthews N., Ellwood M., Woodmansey R., Clark G., Cooper J.D., Tromans A., Grafham D., Skuce C., Pandian R., Andrews R., Harrison E., Kimberley A., Garnett J., Fosker N., Hall R., Garner P., Kelly D., Bird C., Palmer S., Gehring I., Berger A., Dooley C.M., Ersan-Ürün Z., Eser C., Geiger H., Geisler M., Karotki L., Kirn A., Konantz J., Konantz M., Oberländer M., Rudolph-Geiger S., Teucke M., Lanz C., Raddatz G., Osoegawa K., Zhu B., Rapp A., Widaa S., Langford C., Yang F., Schuster S.C., Carter N.P., Harrow J., Ning Z., Herrero J., Searle S.M.J., Enright A., Geisler R., Plasterk R.H.A., Lee C., Westerfield M., de Jong P.J., Zon L.I., Postlethwait J.H., Nüsslein-Volhard C., Hubbard T.J.P., Crollius H.R., Rogers J., Stemple D.L. 2013. The zebrafish reference genome sequence and its relationship to the human genome. Nature. 496:498–503.

32. Hughes L.C., Ortí G., Huang Y., Sun Y., Baldwin C.C., Thompson A.W., Arcila D., Betancur-R. R., Li C., Becker L., Bellora N., Zhao X., Li X., Wang M., Fang C., Xie B., Zhou Z., Huang H., Chen S., Venkatesh B., Shi Q. 2018. Comprehensive phylogeny of ray-finned fishes (Actinopterygii) based on transcriptomic and genomic data. Proc. Natl. Acad. Sci. 115:6249–6254.

33. Jarvis E.D., Mirarab S., Aberer A.J., Li B., Houde P., Li C., Ho S.Y.W., Faircloth B.C., Nabholz B., Howard J.T., Suh A., Weber C.C., da Fonseca R.R., Li J., Zhang F., Li H., Zhou L., Narula N., Liu L., Ganapathy G., Boussau B., Bayzid Md.S., Zavidovych V., Subramanian S., Gabaldón T., Capella-Gutiérrez S., Huerta-Cepas J., Rekepalli B., Munch K., Schierup M., Lindow B., Warren W.C., Ray D., Green R.E., Bruford M.W., Zhan X., Dixon A., Li S., Li N., Huang Y., Derryberry E.P., Bertelsen M.F., Sheldon F.H., Brumfield R.T., Mello C.V., Lovell P.V., Wirthlin M., Schneider M.P.C., Prosdocimi F., Samaniego J.A., Velazquez A.M.V., Alfaro-Núñez A., Campos P.F., Petersen B., Sicheritz-Ponten T., Pas A., Bailey T., Scofield P., Bunce M., Lambert D.M., Zhou Q., Perelman P., Driskell A.C., Shapiro B., Xiong Z., Zeng Y., Liu S., Li Z., Liu B., Wu K., Xiao J., Yinqi X., Zheng Q., Zhang Y., Yang H., Wang J., Smeds L., Rheindt F.E., Braun M., Fjeldsa J., Orlando L., Barker F.K., Jønsson K.A., Johnson W., Koepfli K.-P., O’Brien S., Haussler D., Ryder O.A., Rahbek C., Willerslev E., Graves G.R., Glenn T.C., McCormack J., Burt D., Ellegren H., Alström P., Edwards S.V., Stamatakis A., Mindell D.P., Cracraft J., Braun E.L., Warnow T., Jun W., Gilbert M.T.P., Zhang G. 2014. Whole-genome analyses resolve early branches in the tree of life of modern birds. Science. 346:1320–1331.

34. Kalyaanamoorthy S., Minh B.Q., Wong T.K.F., Von Haeseler A., Jermiin L.S. 2017. ModelFinder: Fast model selection for accurate phylogenetic estimates. Nat. Methods.

35. Katoh K., Standley D.M. 2013. MAFFT multiple sequence alignment software version 7: improvements in performance and usability. Mol. Biol. Evol. 30:772–780.

36. Keilwagen J., Hartung F., Grau J. 2019. GeMoMa: Homology-Based Gene Prediction Utilizing Intron Position Conservation and RNA-seq Data BT - Gene Prediction: Methods and Protocols. In: Kollmar M., editor. New York, NY: Springer New York. p. 161–177.

37. Kim D., Paggi J.M., Park C., Bennett C., Salzberg S.L. 2019. Graph-based genome alignment and genotyping with HISAT2 and HISAT-genotype. Nat. Biotechnol. 37:907–915.

38. Kloosterman W.P., Francioli L.C., Hormozdiari F., Marschall T., Hehir-Kwa J.Y., Abdellaoui A., Lameijer E.-W., Moed M.H., Koval V., Renkens I., van Roosmalen M.J., Arp P., Karssen L.C., Coe B.P., Handsaker R.E., Suchiman E.D., Cuppen E., Thung D.T., McVey M., Wendl M.C., Consortium G. of the N., Uitterlinden A., van Duijn C.M., Swertz M.A., Wijmenga C., van Ommen G.B., Slagboom P.E., Boomsma D.I., Schönhuth A., Eichler E.E., de Bakker P.I.W., Ye K., Guryev V. 2015. Characteristics of de novo structural changes in the human genome. Genome Res. 25:792–801.

39. Kottelat M. 2012. Conspectus cobitidum: an inventory of the loaches of the world (Teleostei: Cypriniformes: Cobitoidei). Raffles Bull. Zool.

40. Krabbenhoft T.J., MacGuigan D.J., Backenstose N.J.C., Waterman H., Lan T., Pelosi J.A., Tan M., Sandve S.R. 2021. Chromosome-Level Genome Assembly of Chinese Sucker (*Myxocyprinus asiaticus*) Reveals Strongly Conserved Synteny Following a Catostomid-Specific Whole-Genome Duplication. Genome Biol. Evol. 13.

41. Kratochwil C.F., Kautt A.F., Nater A., Härer A., Liang Y., Henning F., Meyer A. 2022. An intronic transposon insertion associates with a trans-species color polymorphism in Midas cichlid fishes. Nat. Commun. 13:296.

42. Kuznetsov D., Tegenfeldt F., Manni M., Seppey M., Berkeley M., Kriventseva E.V., Zdobnov E.M. 2023. OrthoDB v11: annotation of orthologs in the widest sampling of organismal diversity. Nucleic Acids Res. 51:D445–D451.

43. Li H. 2013. Aligning sequence reads, clone sequences and assembly contigs with BWA-MEM. ArXiv Prepr. ArXiv13033997.

44. Li H. 2016. Minimap and miniasm: Fast mapping and de novo assembly for noisy long sequences. Bioinformatics.

45. Li H. 2018. Minimap2: pairwise alignment for nucleotide sequences. Bioinformatics. 34:3094–3100.

46. Li H., Durbin R. 2009. Fast and accurate short read alignment with Burrows-Wheeler transform. Bioinformatics.

47. Li H., Handsaker B., Wysoker A., Fennell T., Ruan J., Homer N., Marth G., Abecasis G., Durbin R. 2009. The Sequence Alignment/Map format and SAMtools. Bioinformatics. 25:2078–2079.

48. Li X., Guo B. 2020. Substantially adaptive potential in polyploid cyprinid fishes: evidence from biogeographic, phylogenetic and genomic studies. Proc. R. Soc. B Biol. Sci. 287:20193008.

49. Lin Y.-L., Gokcumen O. 2019. Fine-Scale Characterization of Genomic Structural Variation in the Human Genome Reveals Adaptive and Biomedically Relevant Hotspots. Genome Biol. Evol. 11:1136–1151.

50. Liu D., Hunt M., Tsai I.J. 2018. Inferring synteny between genome assemblies: a systematic evaluation. BMC Bioinformatics. 19:26.

51. Lovell J.T., Sreedasyam A., Schranz M.E., Wilson M., Carlson J.W., Harkess A., Emms D., Goodstein D.M., Schmutz J. 2022. GENESPACE tracks regions of interest and gene copy number variation across multiple genomes. eLife. 11:e78526.

52. Lv Y., Li J., Li Y., Huang Y., Lai Q., Wen Z., Wang J., He Y., Shi J., Huang Z., Jiang Y., de Peer Y.V., Shi Q., Xie B., Wang Y. 2025. Subgenome Partitioning and Polyploid Genome Evolution in the Loach Family Botiidae (Order Cypriniformes). Adv. Sci. n/a:e05411.

53. MacGuigan D.J., Krabbenhoft T.J., Harrington R.C., Wainwright D.K., Backenstose N.J.C., Near T.J. 2023. Lacustrine speciation associated with chromosomal inversion in a lineage of riverine fishes. Evol. Int. J. Org. Evol. 77:1505–1521.

54. Manni M., Berkeley M.R., Seppey M., Zdobnov E.M. 2021. BUSCO: Assessing Genomic Data Quality and Beyond. Curr. Protoc. 1:e323.

55. Martin M. 2011. Cutadapt removes adapter sequences from high-throughput sequencing reads. EMBnet.journal.

56. Mayden R.L., Tang K.L., Conway K.W., Freyhof J., Chamberlain S., Haskins M., Schneider L., Sudkamp M., Wood R.M., Agnew M., Bufalino A., Sulaiman Z., Miya M., Saitoh K., He S. 2007. Phylogenetic relationships of Danio within the order Cypriniformes: a framework for comparative and evolutionary studies of a model species. J. Exp. Zoolog. B Mol. Dev. Evol. 308B:642–654.

57. Mayden R.L., Tang K.L., Wood R.M., Chen W.-J., Agnew M.K., Conway K.W., Yang L., Simons A.M., Bart H.L., M.harris P., Li J., Wang X., Saitoh K., He S., Liu H., Chen Y., Nishida M., Miya M. 2008. Inferring the Tree of Life of the order Cypriniformes, the earth’s most diverse clade of freshwater fishes: Implications of varied taxon and character sampling. J. Syst. Evol. 46:424.

58. Melo B.F., Sidlauskas B.L., Near T.J., Roxo F.F., Ghezelayagh A., Ochoa L.E., Stiassny M.L.J., Arroyave J., Chang J., Faircloth B.C., MacGuigan D.J., Harrington R.C., Benine R.C., Burns M.D., Hoekzema K., Sanches N.C., Maldonado-Ocampo J.A., Castro R.M.C., Foresti F., Alfaro M.E., Oliveira C. 2022. Accelerated Diversification Explains the Exceptional Species Richness of Tropical Characoid Fishes. Syst. Biol. 71:78–92.

59. Meyers J.R. 2018. Zebrafish: Development of a Vertebrate Model Organism. Curr. Protoc. Essent. Lab. Tech. 16:e19.

60. Minh B.Q., Hahn M.W., Lanfear R. 2020a. New Methods to Calculate Concordance Factors for Phylogenomic Datasets. Mol. Biol. Evol. 37:2727–2733.

61. Minh B.Q., Schmidt H.A., Chernomor O., Schrempf D., Woodhams M.D., von Haeseler A., Lanfear R. 2020b. IQ-TREE 2: New Models and Efficient Methods for Phylogenetic Inference in the Genomic Era. Mol. Biol. Evol. 37:1530–1534.

62. Mirarab S., Reaz R., Bayzid Md.S., Zimmermann T., Swenson M.S., Warnow T. 2014. ASTRAL: genome-scale coalescent-based species tree estimation. Bioinformatics. 30:i541–i548.

63. Mirarab S., Rivas-González I., Feng S., Stiller J., Fang Q., Mai U., Hickey G., Chen G., Brajuka N., Fedrigo O., Formenti G., Wolf J.B.W., Howe K., Antunes A., Schierup M.H., Paten B., Jarvis E.D., Zhang G., Braun E.L. 2024. A region of suppressed recombination misleads neoavian phylogenomics. Proc. Natl. Acad. Sci. 121:e2319506121.

64. Near T.J., Eytan R.I., Dornburg A., Kuhn K.L., Moore J.A., Davis M.P., Wainwright P.C., Friedman M., Smith W.L. 2012. Resolution of ray-finned fish phylogeny and timing of diversification. Proc. Natl. Acad. Sci. 109:13698–13703.

65. Nevers Y., Warwick Vesztrocy A., Rossier V., Train C.-M., Altenhoff A., Dessimoz C., Glover N.M. 2025. Quality assessment of gene repertoire annotations with OMArk. Nat. Biotechnol. 43:124–133.

66. Nishimura O., Hara Y., Kuraku S. 2017. GVolante for standardizing completeness assessment of genome and transcriptome assemblies. Bioinformatics.

67. Ou S., Jiang N. 2018. LTR_retriever: A Highly Accurate and Sensitive Program for Identification of Long Terminal Repeat Retrotransposons. Plant Physiol. 176:1410–1422.

68. Parey E., Louis A., Montfort J., Bouchez O., Roques C., Iampietro C., Lluch J., Castinel A., Donnadieu C., Desvignes T., Floi Bucao C., Jouanno E., Wen M., Mejri S., Dirks R., Jansen H., Henkel C., Chen W.-J., Zahm M., Cabau C., Klopp C., Thompson A.W., Robinson-Rechavi M., Braasch I., Lecointre G., Bobe J., Postlethwait J.H., Berthelot C., Roest Crollius H., Guiguen Y. 2023. Genome structures resolve the early diversification of teleost fishes. Science. 379:572–575.

69. Quinlan A.R., Hall I.M. 2010. BEDTools: a flexible suite of utilities for comparing genomic features. Bioinformatics. 26:841–842.

70. Roach M.J., Schmidt S.A., Borneman A.R. 2018. Purge Haplotigs: allelic contig reassignment for third-gen diploid genome assemblies. BMC Bioinformatics. 19:460.

71. Rokas A., Holland P.W.H. 2000. Rare genomic changes as a tool for phylogenetics. Trends Ecol. Evol. 15:454–459.

72. Saitoh K., Sado T., Mayden R.L., Hanzawa N., Nakamura K., Nishida M., Miya M. 2006. Mitogenomic Evolution and Interrelationships of the Cypriniformes (Actinopterygii: Ostariophysi): The First Evidence Toward Resolution of Higher-Level Relationships of the World’s Largest Freshwater Fish Clade Based on 59 Whole Mitogenome Sequences. J. Mol. Evol. 63:826–841.

73. Schlebusch S.A., Trifonov V., Halenková Z., Klianitskaya M., Dedukh D., Herrera A.R., González L.Á., Infantes G.P., Hřibová E., Andjel L., Bartoš O., Pajer P., Tichopád T., Kulik D., Kotusz J., Doležálková M.K., Bohne A., Marta A., Horna P., Reifová R., Guiguen Y., Pačes J., Janko K. 2025. Sex Chromosome Turnover and Structural Interspecific Genome Divergence Shapes Meiotic Outcomes in Hybridizing Cobitis. :2025.04.01.646337.

74. Schultz D.T., Haddock S.H.D., Bredeson J.V., Green R.E., Simakov O., Rokhsar D.S. 2023. Ancient gene linkages support ctenophores as sister to other animals. Nature. 618:110–117.

75. Seppey M., Manni M., Zdobnov E.M. 2019. BUSCO: Assessing Genome Assembly and Annotation Completeness BT - Gene Prediction: Methods and Protocols. In: Kollmar M., editor. New York, NY: Springer New York. p. 227–245.

76. Sodeland M., Jorde P.E., Lien S., Jentoft S., Berg P.R., Grove H., Kent M.P., Arnyasi M., Olsen E.M., Knutsen H. 2016. “Islands of Divergence” in the Atlantic Cod Genome Represent Polymorphic Chromosomal Rearrangements. Genome Biol. Evol. 8:1012–1022.

77. Stanke M., Diekhans M., Baertsch R., Haussler D. 2008. Using native and syntenically mapped cDNA alignments to improve de novo gene finding. Bioinformatics.

78. Steenwyk J.L., King N. 2024a. The promise and pitfalls of synteny in phylogenomics. PLOS Biol. 22:e3002632.

79. Steenwyk J.L., King N. 2024b. The promise and pitfalls of synteny in phylogenomics. PLOS Biol. 22:e3002632.

80. Storer J., Hubley R., Rosen J., Wheeler T.J., Smit A.F. 2021. The Dfam community resource of transposable element families, sequence models, and genome annotations. Mob. DNA. 12:2.

81. Stout C.C., Tan M., Lemmon A.R., Lemmon E.M., Armbruster J.W. 2016. Resolving Cypriniformes relationships using an anchored enrichment approach. BMC Evol. Biol. 16:244.

82. Sturtevant A.H., Dobzhansky Th. 1936. Inversions in the Third Chromosome of Wild Races of Drosophila Pseudoobscura, and Their Use in the Study of the History of the Species. Proc. Natl. Acad. Sci. 22:448–450.

83. Tan M., Armbruster J.W. 2018. Phylogenetic classification of extant genera of fishes of the order Cypriniformes (Teleostei: Ostariophysi). Zootaxa. 4476:6–39.

84. Tan M., Redmond A.K., Dooley H., Nozu R., Sato K., Kuraku S., Koren S., Phillippy A.M., Dove A.D., Read T. 2021. The whale shark genome reveals patterns of vertebrate gene family evolution. eLife. 10:e65394.

85. Tao W., Yang L., Mayden R.L., He S. 2019. Phylogenetic relationships of Cypriniformes and plasticity of pharyngeal teeth in the adaptive radiation of cyprinids. Sci. China Life Sci. 62:553–565.

86. Tonini J., Moore A., Stern D., Shcheglovitova M., Ortí G. 2015. Concatenation and Species Tree Methods Exhibit Statistically Indistinguishable Accuracy under a Range of Simulated Conditions. PLoS Curr. 7.

87. Townsend J.P. 2007. Profiling Phylogenetic Informativeness. Syst. Biol. 56:222–231.

88. Vaser R., Sović I., Nagarajan N., Šikić M. 2017. Fast and accurate de novo genome assembly from long uncorrected reads. Genome Res.

89. Walker B.J., Abeel T., Shea T., Priest M., Abouelliel A., Sakthikumar S., Cuomo C.A., Zeng Q., Wortman J., Young S.K., Earl A.M. 2014. Pilon: An integrated tool for comprehensive microbial variant detection and genome assembly improvement. PLoS ONE.

90. Wang Y., Tang H., DeBarry J.D., Tan X., Li J., Wang X., Lee T., Jin H., Marler B., Guo H., Kissinger J.C., Paterson A.H. 2012. MCScanX: a toolkit for detection and evolutionary analysis of gene synteny and collinearity. Nucleic Acids Res. 40:e49–e49.

91. Waterhouse R.M., Seppey M., Simão F.A., Manni M., Ioannidis P., Klioutchnikov G., Kriventseva E.V., Zdobnov E.M. 2018. BUSCO Applications from Quality Assessments to Gene Prediction and Phylogenomics. Mol. Biol. Evol. 35:543–548.

92. Whitfield J.B., Lockhart P.J. 2007. Deciphering ancient rapid radiations. Trends Ecol. Evol. 22:258–265.

93. Wilson C.A., High S.K., McCluskey B.M., Amores A., Yan Y., Titus T.A., Anderson J.L., Batzel P., Carvan M.J. III, Schartl M., Postlethwait J.H. 2014. Wild Sex in Zebrafish: Loss of the Natural Sex Determinant in Domesticated Strains. Genetics. 198:1291–1308.

94. Wood D.E., Lu J., Langmead B. 2019. Improved metagenomic analysis with Kraken 2. Genome Biol.

95. Yu H., Li Y., Han W., Bao L., Liu F., Ma Y., Pu Z., Zeng Q., Zhang L., Bao Z., Wang S. 2024. Pan-evolutionary and regulatory genome architecture delineated by an integrated macro- and microsynteny approach. Nat. Protoc. 19:1623–1678.

96. Zhang C., Scornavacca C., Molloy E.K., Mirarab S. 2020. ASTRAL-Pro: Quartet-Based Species-Tree Inference despite Paralogy. Mol. Biol. Evol. 37:3292–3307.

97. Zhang F., Ding Y., Zhu C., Zhou X., Orr M.C., Scheu S., Luan Y. 2019. Phylogenomics from low-coverage whole-genome sequencing. Methods Ecol. Evol. 10:507–517.

98. Zhao T., Zwaenepoel A., Xue J.-Y., Kao S.-M., Li Z., Schranz M.E., Van de Peer Y. 2021. Whole-genome microsynteny-based phylogeny of angiosperms. Nat. Commun. 12:3498.

